# Regulation of gene drive expression increases invasive potential and mitigates resistance

**DOI:** 10.1101/360339

**Authors:** Andrew Hammond, Xenia Karlsson, Ioanna Morianou, Kyros Kyrou, Andrea Beaghton, Matthew Gribble, Nace Kranjc, Roberto Galizi, Austin Burt, Andrea Crisanti, Tony Nolan

## Abstract

CRISPR-Cas9 nuclease-based gene drives rely on inducing chromosomal breaks in the germline that are repaired in ways that lead to a biased inheritance of the drive. Gene drives designed to impair female fertility can suppress populations of the mosquito vector of malaria. However, strong unintended fitness costs, due to ectopic nuclease expression, and high levels of resistant mutations, limited the potential of the first generation of gene drives to spread.

Here we show that changes to regulatory sequences in the drive element, designed to contain nuclease expression to the germline, confer improved fecundity over previous versions and generate drastically lower rates of target site resistance. We employed a genetic screen to show that this effect is explained by reduced rates of end-joining repair of DNA breaks at the target site caused by deposited nuclease in the embryo.

Highlighting the impact of deposited Cas9, many of the mutations arising from this source of nuclease activity in the embryo are heritable, thereby having the potential to generate resistant target sites that reduce the penetrance of the gene drive.

Finally, in cage invasion experiments these gene drives show improved invasion dynamics compared to first generation drives, resulting in greater than 90% suppression of the reproductive output and a delay in the emergence of target site resistance, even at a resistance-prone target sequence. We shed light on the dynamics of generation and selection of resistant alleles in a population by tracking, longitudinally, the frequency of resistant alleles in the face of an invading gene drive. Our results illustrate important considerations for future gene drive design and should expedite the development of gene drives robust to resistance.

## Introduction

### Gene drives and malaria control

Gene drives are genetic elements that are capable of biasing their own inheritance, allowing their autonomous spread in a population, even from a very low initial frequency. Coupling a trait of interest to a drive element is a way of deliberately modifying a population, potentially in a very short timeframe. In the case of the mosquito vector of malaria, gene drives have been proposed to spread traits that either interfere with the mosquito’s capacity to reproduce, or its capacity to transmit the malaria parasite.

We know that vector control is effective in controlling malaria – the malaria burden was halved in the period 2000-2015 and the vast majority of this gain was achieved through targeting the vector with conventional insecticide-based approaches (bednets and residual spraying) [1]. However, resistance to insecticides is stalling these gains [2]. Gene drive is a technology with the potential to augment and complement existing control approaches in a self-sustaining way.

### Endonuclease-based homing gene drives

Gene drives based on site-specific endonucleases were first proposed over 15 years ago [3] and recent advances in CRISPR technology have led to several demonstrations that this endonuclease, which is easy to reprogram to recognise a genomic site of choice, can be repurposed as a gene drive [4, 5].

The premise is that the endonuclease is sufficiently specific to recognise a DNA target sequence within a region of interest and the gene encoding the endonuclease is inserted within this target sequence on the chromosome, thereby rendering it immune to further cleavage. When a chromosome containing the endonuclease is paired with a chromosome containing the wild type target site, the site is cleaved to create a double stranded break (DSB) that can be repaired, either through simple ‘cut and shut’ non-homologous end-joining (NHEJ) or through homology-directed repair (HDR). HDR involves strand invasion from the broken strand into regions of immediate homology on the intact chromosome, and synthesis across the intervening region to repair the gap. In the arrangement described this can lead to copying of the endonuclease, and its associated allele, from one chromosome to another in a process referred to as ‘homing’. If homing takes place in the germline then inheritance of the gene drive is biased because the majority of sperm or eggs produced in the germline will inherit the drive, on either the original gene drive-carrying chromosome or the newly converted chromosome. The rate of spread of this type of gene drive is thus a product of the rate of germline nuclease activity and the probability with which double stranded breaks are repaired by the host cell using HDR rather than end-joining repair.

### Previous limitations of CRISPR-based homing gene drives

We and others have tested several gene drives designed to suppress or modify mosquito populations [6-8]. With any gene drive the force of selection for resistance to the drive will be proportional to the fitness cost imposed by the drive. In the case of population suppression approaches, this selection falls on the mosquito. We previously built several CRISPR-based gene drives designed to achieve population suppression through disrupting female fertility genes and demonstrated their spread throughout caged populations of the malaria mosquito, *Anopheles gambiae* [7]. However, the drive was eventually replaced in the population by resistant mutations generated by end-joining repair in the fraction of cleaved chromosomes that were not modified by homing [9]. To ultimately replace the drive in a population, these mutations must also encode a functional copy of the target gene so that they restore fertility to females.

In determining the propensity for target site resistance to arise, functional constraint at the target site is paramount to determine the degree of variation that can be tolerated there. The target site sequence in the first iteration of a population suppression gene drives is poorly conserved [7], suggesting little functional constraint, and therefore is particularly prone to accommodating resistant alleles that restore function to the target gene (‘r1’ alleles, as opposed to ‘r2’ alleles which are resistant to cleavage but do not restore function to the target gene) (Supplementary Figure 1) [9, 10]. The importance of choosing functionally constrained sites was shown when a gene drive targeting a highly conserved target sequence in the female-specific isoform of a sex determination gene was able to crash caged populations without selecting for resistant mutations of the r1 class [8]. Notwithstanding this success, it is unlikely that any single site will be completely ‘resistance-proof’. This is true for any suppressive technology, from antibiotics to insecticides, and measures to prolong the durability of these interventions need to be considered at inception.

In addition to multiplexing gene drives to recognise more than one target site [3, 4, 11], akin to combination therapy with antibiotics, it is necessary to reduce the relative contribution of the error-prone end-joining repair pathway, over HDR, since this can serve to increase the range and complexity of potential resistant alleles on which selection could act.

Cleavage by maternally deposited nuclease in the embryo is believed to be the major source of end-joining mutations in current gene drive designs [6, 9, 12]. We have previously observed high levels of end-joining repair in the embryo following strong maternal deposition using the *vas2* promoter. This is likely a consequence of germline expression that persists through oocyte development leading to perduring Cas9 transcript and/or protein in the newly fertilised embryo. Additionally, using the same promoter, unrestricted activity of the endonuclease outside of the mosquito germline caused a strong unintended fitness cost in females harbouring a single copy of the gene drive, due to partial conversion of the soma to the homozygosity for a null allele. These fitness effects retarded the invasive potential of the drive for two reasons: 1-reduced bias in inheritance each generation due to low fecundity of drive-carrying (heterozygous) females; 2-increased selection pressure for resistant mutations [9].

Given the limitations of the promoter in previous gene drive constructs we decided to test a suite of novel germline promoters for their ability to restrict nuclease activity to the germline. We employed genetic screens to get a quantitative and qualitative determination of the mutations arising from the different gene drive constructs that, importantly, allowed us to determine where the mutations were generated. We then employed the best performing constructs in a cage invasion assay to determine how our single generation estimates of homing performance translated to our mathematical models of penetrance and population suppression over time, while tracking the emergence of resistant alleles.

## Results and Discussion

### Choice of germline promoters

To find alternatives to the *vas2* promoter, we investigated three mosquito genes, *AGAP006241* (*zero population growth, zpg*), *AGAP006098* (*nanos, nos*) and *AGAP007365* (*exuperantia, exu*), that are expressed in the germline of male and female *Anopheles gambiae* [13] and may show reduced somatic expression or deposition into the embryo.

In *Drosophila* the gene *zpg* is specifically expressed in the male and female germline, where it mediates the formation of gap junctions between the developing germline and cyst cells [14]. The mosquito ortholog of *zpg* appears to be functionally conserved as it is essential for both male and female gonad development [15]. *Exu* and *nos* are maternal effect genes in *Drosophila* that are transcribed in the oocyte and deposited into the early embryo [16-18]. Crucially, deposited *nos* and *zpg* mRNA concentrate at the germ plasm due to regulatory signals present on the untranslated regions, which also further restrict translation of maternal mRNAs to the germline [14, 19]. The promoter region of *exuperantia* has been validated in *Drosophila* [20] and in the tiger mosquito, *Aedes aegypti*, has been used to drive robust expression in both male and female germlines and has recently been used to control expression of Cas9 in a split gene drive system in this mosquito [21, 22]. In contrast to *zpg* and *exu*, several reports have suggested that *nos* is specific to the female germline in mosquitoes, however promoter fusions in *Aedes aegypti* and *Anopheles gambiae* led to low level expression in males perhaps due to incomplete recapitulation of endogenous gene expression [23-25].

### Generation of new gene drives designed to restrict spatiotemporal expression to the mosquito germline

To determine the effect of transcriptional control of gene drive activity, we tested the new constructs at a previously validated female fertility locus that was prone to resistance [9], in order to quantify any reduction in the creation or selection of resistant mutations that resulted from reducing embryonic end-joining and improving female fertility, respectively.

The target site in question resides within the gene *AGAP007280* (Fig. 1A), an ortholog of the *Drosophila* gene *nudel* required in the follicle cells to define polarity of the eggshell [26, 27]. Null mutations at this target site cause a recessive female sterile phenotype. The active gene drive cassette (*CRISPR*^*h*^) used previously to target *AGAP007280* [7] was modified to contain Cas9 under control regulatory sequences flanking upstream and downstream of either *zpg, nos* or *exu* genes. The *CRISPR*^*h*^ constructs were inserted within the *AGAP007280* target site, by recombinase-mediated cassette exchange, as previously [7] (Fig. 1B). The new gene drive strains were named *nos-CRISPR*^*h*^, *zpg-CRISPR*^*h*^, *exu-CRISPR*^*h*^, respectively.

**Figure 1.**
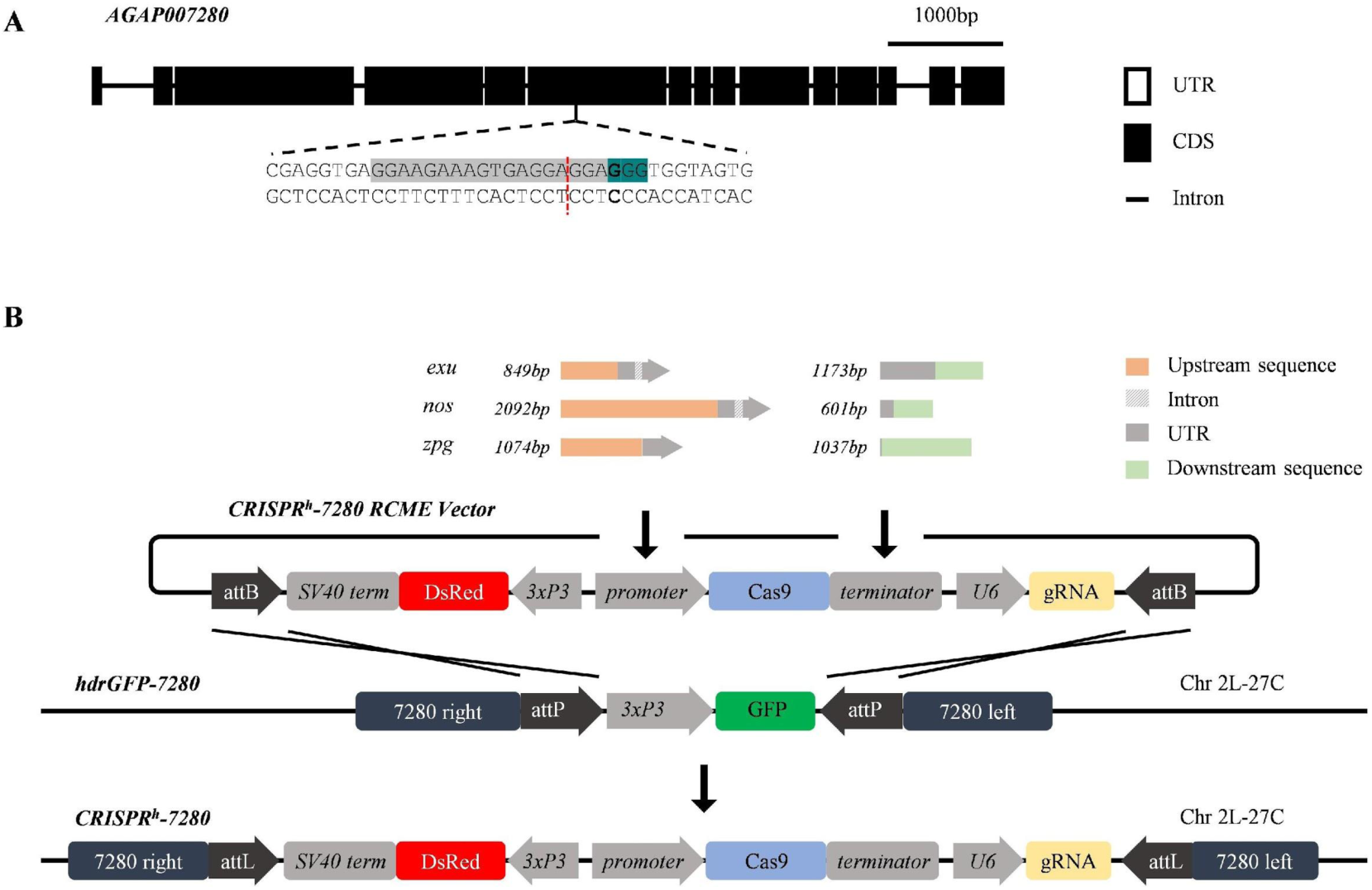
Target site and design of new *CRISPR*^*h*^ gene drives designed to express Cas9 under regulation of *zpg, nos* and *exu* germline promoters. (**a**) The haplosufficient female fertility gene, *AGAP007280*, and its target site in exon 6 (highlighted in grey), showing the protospacer-adjacent motif (highlighted in teal) and cleavage site (red dashed line). (**b**) *CRISPR*^*h*^ alleles were inserted at the target in *AGAP007280* using ϕC31-recombinase mediated cassette exchange (RCME). Each *CRISPR*^*h*^ RCME vector was designed to contain *Cas9* under transcriptional control of the *nos, zpg* or *exu* germline promoter and terminator, a gRNA targeted to *AGAP007280* under the control of the ubiquitous *U6* PolIII promoter, and a *3xP3∷DsRed* marker.

### *Zpg* and *nos* promoters drive high levels of homing in the germline and vary in magnitude of maternal effect

Germline gene drive activity that leads to homing is expected to cause super-Mendelian inheritance of the drive. However, maternally-deposited Cas9 has the potential to cause resistant mutations in the embryo that may reduce the rate of homing during gamete formation (if the mutations occur in germline precursor cells) or reduce fertility (due to somatic mosaicism of null mutations in the target fertility gene) [6, 9, 10, 12, 28].

Assays were performed to measure the transmission rates and fertility costs associated with each of three drives (Fig. 2). Individuals heterozygous for the gene drive were crossed to wild type and their progeny scored for the presence of the DsRed marker gene linked to the construct. To test the magnitude of any parental effect, gene drive carriers were separated according to whether they received their gene drive allele paternally or maternally. Data from our previously generated gene drive constructs under control of the vasa promoter (*vas2-CRISPR*^*h*^) served as a benchmark [7].

**Figure 2.**
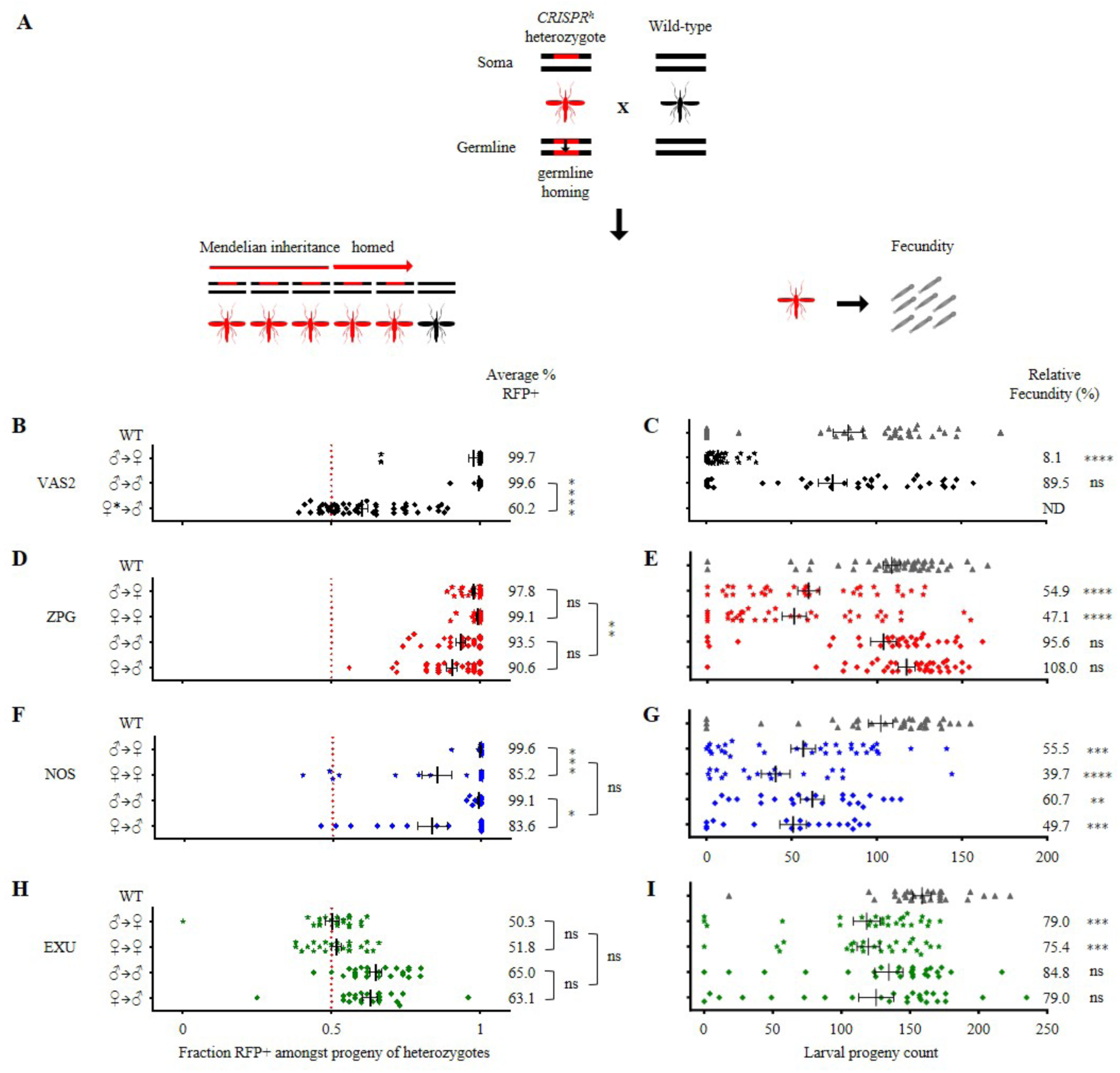
Comparison of fecundity and homing rates in mosquitoes containing Cas9-based gene drives under regulation of *zpg, nos* and *exu* germline promoters. Phenotypic assays were performed to measure fertility and transmission rates for each of three drives. The larval output was determined for individual drive heterozygotes crossed to wild type (left, B,D,F,H), and their progeny scored for the presence of DsRed linked to the construct (right,C,E,G,I). Average transmission rates are shown on the right of panels B, D, F, H and relative fecundity to wild type is depicted on the right of panels C, E, G, I (Kruskal-Wallis test, ****: *p* < 0.0001, ***: *p* < 0.001,**: *p* < 0.01, ns: non-significant). Males and females were further separated by whether they had inherited the *CRISPR*^*h*^ construct from either a male or female parent. For example, ♂→♀ denotes progeny and transmission rates of a heterozygous *CRISPR*^*h*^ female that had inherited the drive allele from a heterozygous *CRISPR*^*h*^ male. High levels of homing were observed in the germline of *zpg-CRISPR*^*h*^ and *nos-CRISPR*^*h*^ males and females, however the *exu* promoter generated only moderate levels of homing in the germline of males but not females. The significant maternal effect upon homing performance in offspring previously seen for *vas2-CRISPR*^*h*^ (Hammond et al. 2017, shown here) was also observed in *nos-CRISPR*^*h*^, but not for *zpg-CRISPR*^*h*^ (Mann-Whitney, ****: *p* < 0.0001, ***: *p* < 0.001, *: *p* < 0.05, ns: non-significant). Counts of hatched larvae for the individual crosses revealed improvements in the fertility of heterozygous females containing *CRISPR*^*h*^ alleles based upon *zpg, nos* and *exu* promoters compared to the *vas2* promoter. Phenotypic assays were performed on G2 and G3 for *zpg*, G3 and G4 for *nos*, and ∼G15 for *exu*. ♀* denotes *vas2-CRISPR*^*h*^ females that were heterozygous for a resistance (r1) allele, these were used because heterozygous *vas2-CRISPR*^*h*^ females are usually sterile.

When the gene drive was received paternally *zpg-CRISPR*^*h*^ transmission rates of 93.5% (±1.5% s.e.m.) in males and 97.8% (±0.6% s.e.m.) in females were observed, falling only slightly below the *vas2-CRISPR*^*h*^ rates in the equivalent cross (99.6% ±0.3% s.e.m. in males and 97.7% ±1.6% s.e.m. in females) (Fig. 2D, Fig. 2B). The paternally-received *nos-CRISPR*^*h*^ construct showed comparable rates to *vas2-CRISPR*^*h*^, with 99.6% (±0.3% s.e.m.) inheritance in females and 99.1% (±0.3% s.e.m.) inheritance in males. In contrast, paternally-received *exu-CRISPR*^*h*^ showed only modest homing rates in males (65.0% ±2.0% s.e.m. transmission rate) and no homing in females (Fig. 2H).

When the gene drive was received maternally, the *vas2-CRISPR*^*h*^ constructs showed a large reduction in homing in males (60.2% ±1.9% s.e.m. inheritance rate, equivalent to homing of 20.2% of non-drive chromosomes) compared to those that received the drive paternally (99.2% homing of non-drive chromosomes) (Fig. 2B). On the contrary, for the *zpg*-*CRISPR*^*h*^ construct, we saw minimum transmission rates of 90.6% (±1.8% s.e.m.) in males and a higher rate of 99.1% in females (±0.4% s.e.m.) (p<0.01), yet no significant maternal or paternal effect (Fig. 2D). With the *nos-CRISPR*^*h*^ construct, though homing rates were still high, we saw a strong maternal effect leading to reduction in homing performance in both males (83.6% ±5.0% s.e.m. inheritance when inherited maternally vs 99.1% ±0.4% s.e.m. when paternally inherited, p<0.05) and females (85.2% ±5.0% s.e.m. vs 99.6% ±0.3% s.e.m., respectively, p<0.001) (Fig. 2F). This suggests that Cas9 from the *nos-CRISPR*^*h*^ construct is maternally deposited, though to a lesser extent than the *vas2-CRISPR*^*h*^ constructs, and active in a way that leads to the formation of alleles in the zygotic germline that are resistant to homing in the next generation. Finally, maternally-inherited *exu-CRISPR*^*h*^ constructs showed modest homing rates in males (63.1% ±2.4% s.e.m. inheritance), unaffected by parental effects, though the absence of gene drive activity in females (51.8% ±1.7% s.e.m. inheritance) may imply that there was simply insufficient nuclease expression in the female germline to produce a maternal effect (Fig. 2H).

### New gene drive constructs confer significantly less fecundity costs in carrier females than previous constructs

Fecundity assays were performed to quantify the number of viable progeny (measured as larval output) in individual crosses of drive heterozygotes mated to wild type (Fig. 2). Our previously published data for *vas2-CRISPR*^*h*^ females showed vastly impaired fecundity (8% fecund, relative to wild type) in females heterozygous for this construct at the *AGAP007280* locus, despite this gene showing full haplosufficiency for this phenotype (i.e. individuals heterozygous for a simple null allele show normal fecundity)[7]. The most parsimonious explanation for this fitness effect is the (partial) conversion of the soma to the null phenotype, due to two sources: somatic ‘leakiness’ of the putative germline promoter driving the Cas9 nuclease; the stochastic distribution (mosaicism) across the soma and germline of nuclease parentally deposited into the embryo.

Both the *zpg-CRISPR*^*h*^ and the *nos-CRISPR*^*h*^ drives showed a general marked improvement in relative female fecundity over the previous *vas2*-driven construct, with maximal fecundities (compared to wild type) of 54.9% and 55.5%, respectively. Although we observed significant maternal deposition from the *nos-CRISPR*^*h*^ construct, the reduction in fecundity of females that received the gene drive maternally was non-significant, compared to those that received the drive paternally (39.7% ±8.5% s.e.m. vs 55.5% ±7.3 s.e.m. larval output respectively) (Fig. 2G). For this construct we also noted a general reduction in male fecundity, regardless of paternal source of the gene drive allele. For *zpg-CRISPR*^*h*^ we observed no reduction in male fecundity and the reduction in female fecundity was not subject to any maternal effect, consistent with the absence of maternal effect on homing for the same construct. Thus, the most likely explanation for the improved performance of the *zpg-CRISPR*^*h*^ gene drive is the lack of a maternal effect, though the incomplete restoration of full fecundity in heterozygote females suggests there may still be some somatic leakiness.

### Embryonic and germline rates of end-joining induced by gene drives containing *zpg* and *nos* promoters

Given its importance in the generation of resistant alleles, we designed a genetic screen to quantify the magnitude of parental nuclease deposition from each construct (Fig 3A). In this screen the wild-type target allele in the embryo, balanced against a pre-existing r1 allele (a 6bp GAGGAG deletion) is only exposed to a paternal or maternal dose of the nuclease in the absence of a genetically encoded drive construct. We performed amplicon sequencing of the relevant region of the *AGAP007280* target locus to sample and quantify the diversity of alleles at the target site. Two replicates were performed for each cross. In the absence of any Cas9 activity an equal ratio of the wild-type target site and the original r1 allele is expected among the chromosomes amplified. Novel indels arising at the target sequence are indicative of nuclease activity in the zygote as a result of parental deposition. The original r1 allele may also be over-represented among the F2 in this assay due to simple stochastic variation in its Mendelian inheritance or due to deposited Cas9 activity, either as a result of *de novo* generation through end-joining repair, or as a result of gene conversion through homing of the inherited r1 allele. Taking only the most conservative signal of embryonic end-joining (indels unique from the inherited r1 allele sequence), *vas2-CRISPR*^*h*^ generated high levels of maternally deposited Cas9 activity – affecting 70% of all nuclease-sensitive alleles (Figure 3B and Supp Figure 3). In the offspring of *nos-CRISPR*^*h*^ females the end-joining rates from maternal deposition were substantially lower (10.5% of nuclease-sensitive alleles), with no significant paternal deposition. For the *zpg-CRISPR*^*h*^ line, we found no significant signal of maternal or paternal deposition (Supp Figure 3).

**Figure 3.**
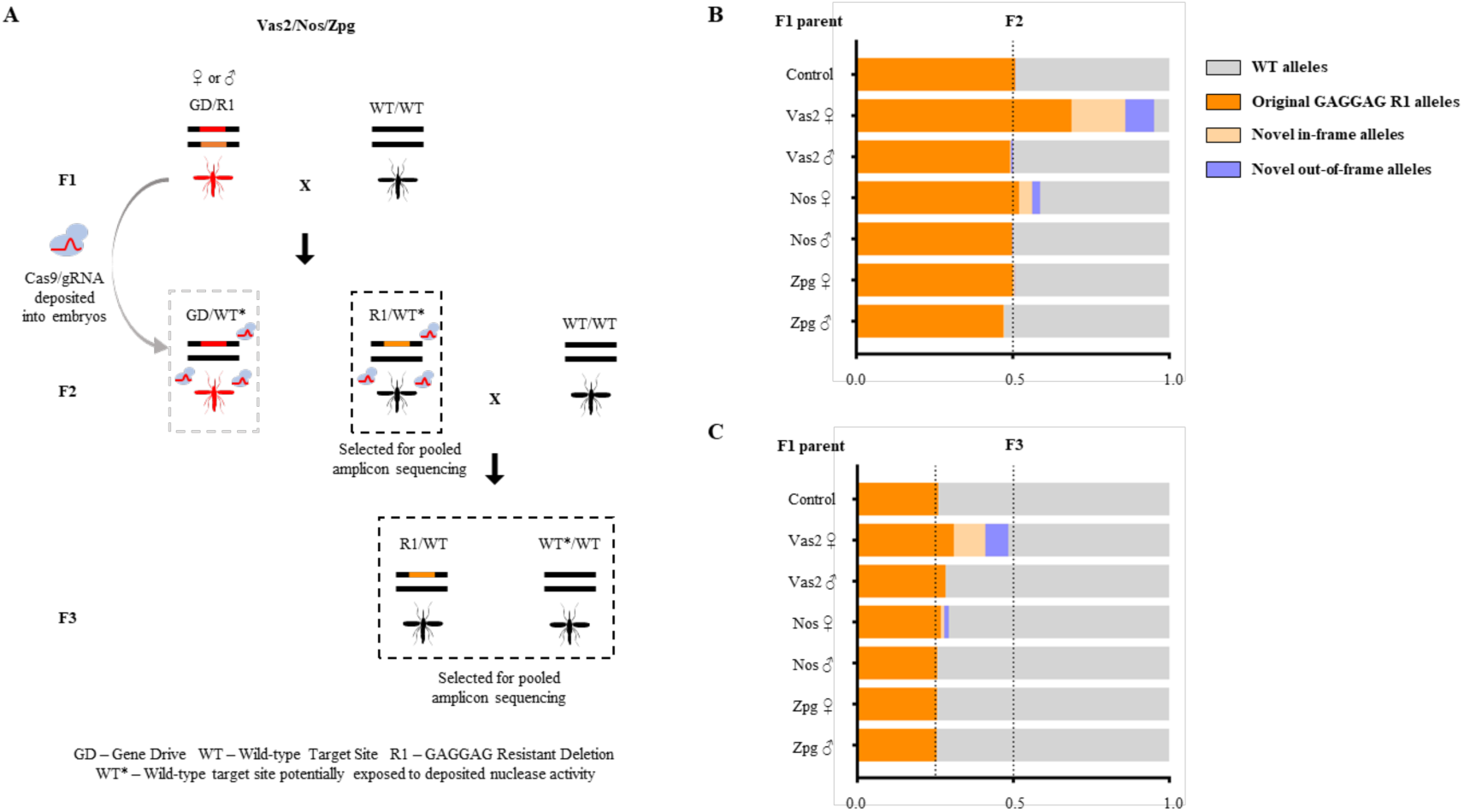
Heritability and rates of formation of mutations caused by parentally deposited nuclease from different gene drive constructs. **(A)** *zpg-CRISPR*^*h*^, *nos-CRISPR*^*h*^ or *vas2-CRISPR*^*h*^ were crossed to a homozygous resistant strain (r1 203-GAGGAG) to generate F1 heterozygotes containing both a gene drive and resistant allele (GD/r1). F1 heterozygotes and r1/r1 homozygotes (control) were crossed to wild type and their non-drive F2 progeny analysed by pooled amplicon sequencing across the target site in *AGAP007280*. **(B)** Amplicon sequencing results from F2 individuals with genotype wt/R1 (selected by absence of dsRED-linked gene drive allele) whose only source of Cas9 nuclease would be from parental deposition into the embryo. In the absence of any deposited source of nuclease only the original r1 allele and the wt allele are expected, at a ratio of 1:1 (**C**) Amplicon sequencing of F3 progeny deriving from the F2 crossed to wild type. Mendelian inheritance of mutations present in the F2 would be expected to lead to a 2-fold reduction in their frequency between the F2 and F3.

Mutant alleles generated in the embryo, if included in the germline tissue, will be resistant to subsequent homing during the production of gametes in the adult, reducing rates of drive transmission. Indeed, the gradation in terms of maternal contribution to end joining in the embryo (*vas2-CRISPR*^*h*^ > *nos-CRISPR*^*h*^ > *zpg-CRISPR*^*h*^) mirrors the magnitude of the maternal effect observed in reducing drive transmission for each construct (**Figure 2**). Consistent with this, the presence in the F3 of novel end-joining mutations that were created in the F2 confirms that deposited nuclease affected also the germline tissue (**Figure 3C**). However, the frequency of resistant alleles generated in the F2 was approximately halved in the F3, consistent with their simple Mendelian inheritance. This indicates that any contribution of deposited nuclease to homing of these alleles in the germline is at best minimal. This is in contrast to reports in *Drosophila* where use of the *nanos* promoter can lead to ‘shadow drive’ in which perduring maternally deposited Cas9 can cause in homing in the germline even in the absence of genetically encoded Cas9 [29, 30].

Given the improved characteristics of the *zpg-CRISPR*^*h*^ construct in terms of fecundity and reduced embryonic end-joining we also investigated its propensity to generate end-joining mutants in the germline. We designed a genetic cross that allowed us to enrich and capture those chromosomes (∼2-3%) that were subjected to the nuclease activity of the *zpg-CRISPR*^*h*^ gene drive in the adult germline but were not ‘homed’ (Fig 4). Interestingly from males (n=240) we noticed a much greater heterogeneity of alleles (82 mutated chromosomes covered by 18 distinct alleles) among those generated from end-joining repair than we did in females (n=219) where the vast majority (88%) of non-homed chromosomes were mutated but these comprised just 4 unique alleles. The apparent differences between the male and female germlines may reflect differences in the timing of nuclease activity and chromosomal repair, and may suggest that early repair events in the female germline, when only a few germline stem cells are present, leads to a clustering of repair events among the eventual gametes.

**Figure 4.**
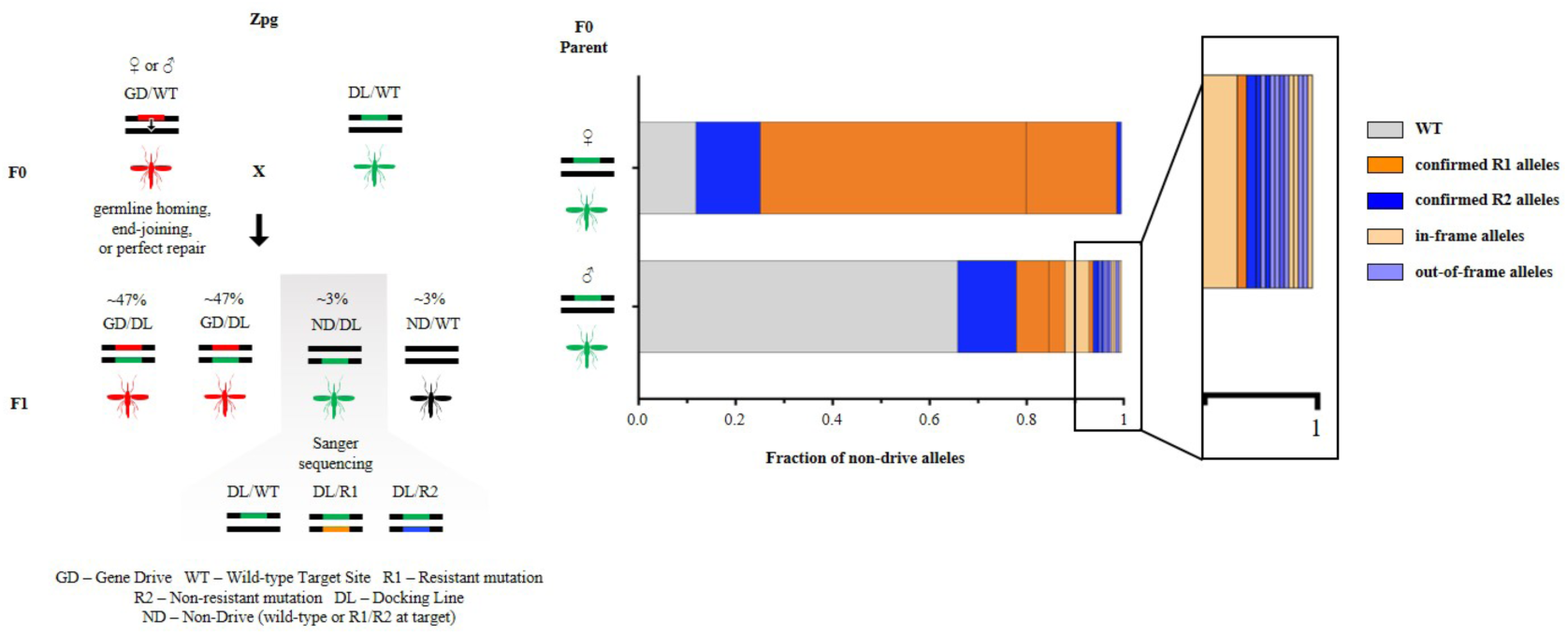
The rate and outcome of meiotic end-joining (EJ) differs in the male and female germline. F0 females (n>500) or males (n>200) heterozygous for the gene drive (linked to RFP+) (GD) were crossed to GFP+ docking line (DL). The rare fraction (∼3%) of F1 progeny that had inherited a DL but not a GD allele were selected and their non-drive allele (ND), that had been meiotically exposed to nuclease activity, was sequenced to reveal the frequency and nature of EJ mutations taking place during meiosis (offspring from females n=219; offspring from males n=240). The frequency of all WT unmodified alleles is shown in grey. Mutant alleles individually exceeding 1% frequency were identified as r1 (orange) or r2 (blue). Mutations present at less than 1% frequency were grouped together as in-frame (light orange) or out-of-frame (light blue).

### *zpg-CRISPR*^*h*^ spreads to close to fixation in caged releases and exerts a large reproductive load on the population

Given its improved fecundity and its lower mutagenic activity we investigated the potential for the *zpg-CRISPR*^*h*^ gene drive to spread throughout naïve mosquito populations. Two replicate cages were initiated with either 10% or 50% starting frequency of drive heterozygotes, and monitored for 16 generations, which included pooled sequencing of the target locus at various generations. The drive spread rapidly in all four trials, to more than 97% of the population, achieving maximal frequency in just 4-10 generations (**Figure 5a**). In each trial, the drive sustained more than 95% frequency for at least 3 generations before its spread was reversed by the gradual selection of drive-resistant alleles. Notably, we observed similar dynamics of spread, in terms of rate of increase and duration of maximal frequency, whether the gene drive was released at 50% or 10%, demonstrating that initial release frequency has little impact (providing it is not stochastically lost initially) upon the potential to spread.

**Figure 5.**
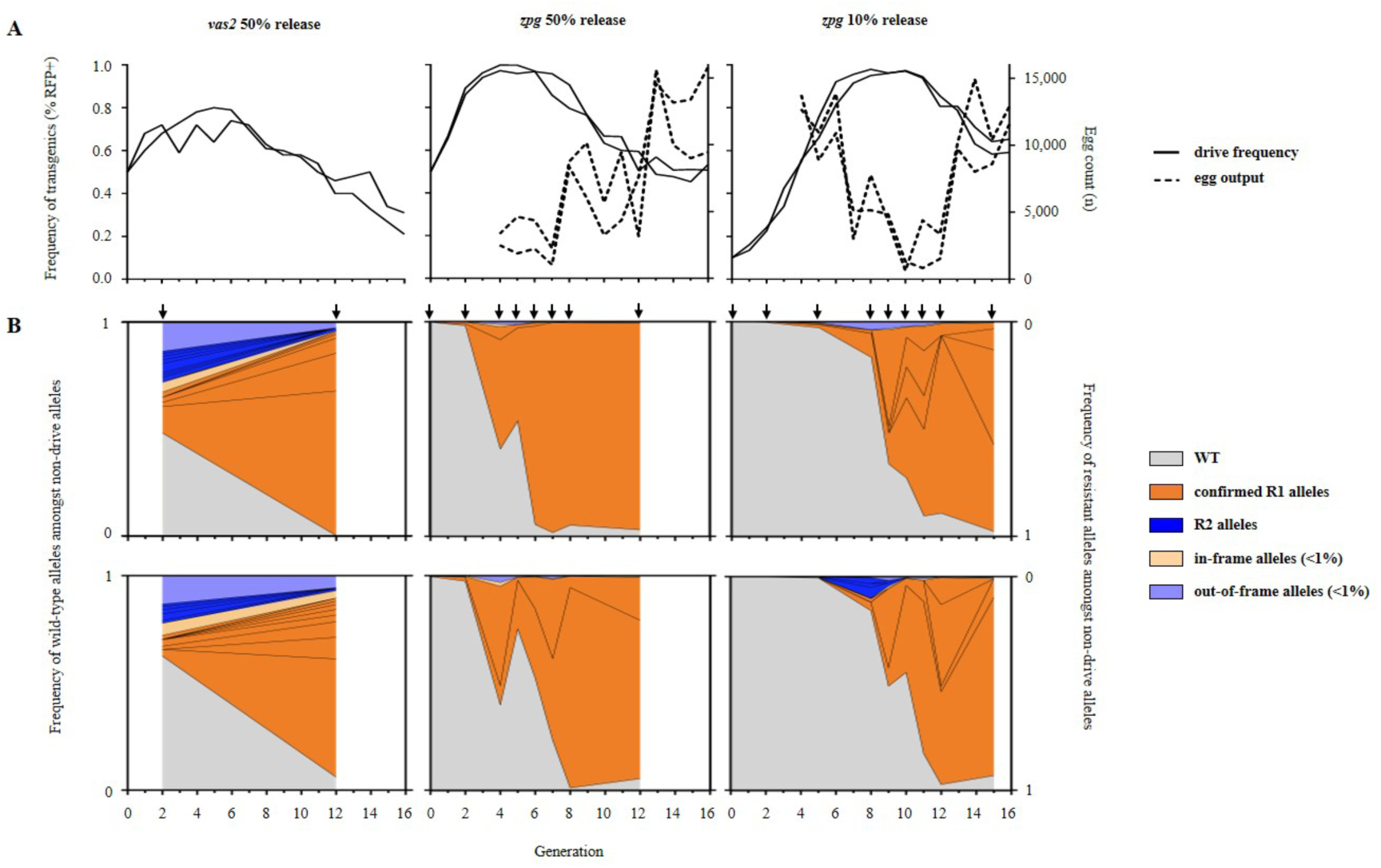
*zpg-CRISPR*^*h*^ rates of spread through caged populations, reproductive load and real-time profiling of resistance. *zpg-CRISPR*^*h*^/*+* were released into replicate caged WT populations at 10% (right) or 50% (middle) *CRISPR*^*h*^/*+* initial frequency. **(a)** The frequency of drive-modified mosquitoes (solid line) was recorded each generation by screening larval progeny for presence of DsRed linked to the *CRISPR*^*h*^ allele. Entire egg output per generation was also recorded (dashed line). *zpg-CRISPR*^*h*^ drive spread was compared to previous results for *vas2-CRISPR*^*h*^ (left panel) from Hammond *et al*. (2017). **(b)** The nature and frequency of non-drive alleles was determined for several generations (arrows) of the caged population experiments by amplicon sequencing across the target site at *AGAP007280* in pooled samples. The frequency of all WT alleles is shown in grey. Mutant alleles individually exceeding 1% frequency in any generation were labelled as r1 (orange) or r2 (blue). Mutations that were individually present at less than 1% frequency were grouped together as in-frame (light orange) or out-of-frame (light blue).

The rates of invasion observed here represent a significant improvement over the first generation gene drive (*vas2-CRISPR*^*h*^) targeted to the exact same target sequence (included for comparison in **Figure 5a**), where the spread of the drive was markedly slower and declined before it reached 80% frequency in the population [7].

Our population suppression gene drive is designed to exert a reproductive load on the population by targeting a gene that is essential for viable egg production. To investigate the magnitude of this load, we counted (from generation 4) the number of viable eggs produced each generation to measure the level of population suppression. Egg production was suppressed by an average of 92% (compared to maximal output of the cage) in each cage at or shortly after the peak in drive frequency, representing a reduction from more than 15,000 eggs to under 1,200 eggs. Since we set a fixed carrying capacity for the adult population in the cage, selecting 600 larvae each generation, the reduction in egg output, although severe, was insufficient to suppress the population below this carrying capacity. In practice, in the wild, the effect of these levels of reproductive suppression will depend very much on the strength of density-dependent mortality that is occurring in the relevant mosquito larval habitats [31, 32].

### The creation and selection of resistant mutations is delayed in cage invasion experiments when using the *zpg* promoter

Both r1 mutations and r2 mutations can retard the invasiveness of a gene drive, yet r1 mutations are more problematic since they are likely to be strongly selected due to the relative positive fitness they confer by restoring function to the target gene. To determine the rate of accumulation of these mutations during the invasion experiment we sequenced across the target locus in samples of pooled individuals from intermittent generations that spanned the rise and fall in frequency of the gene drive (**Figure 5b**).

For the *zpg-CRISPR*^*h*^ constructs, which have minimal fitness effects in carrier females (containing one drive allele and one wild type allele), resistance did not show obvious signs of selection (judged by large increases in frequency) until more than 90% of individuals were positive for the gene drive. In contrast, our previous data for the *vas2*-driven constructs showed that resistance was selected much earlier, being already present at high frequency in the 2^nd^ generation when the frequency of drive-positive individuals was less than 70%. This is likely due to a combination of factors: the somatic leakiness of the *vas2* promoter resulting in a significant fitness cost in carrier females and thus a relative fitness benefit to the r1 allele at a much earlier stage in the invasion, even when drive homozygotes are still rare; the high initial frequency of r2 alleles, resulting in many individuals that are null (either homozygous for the r2 allele or contain one r2 allele and one drive allele) for the target gene therefore meaning that any non-null allele has a stronger relative fitness advantage.

Not only does the *zpg* promoter delay the onset of drive resistance, it also reduces the range of resistant alleles created (**Fig 5b**). In the 50% release cages, only 2 mutant alleles were detected above threshold frequency (>1% of non-drive alleles) both of which had been previously confirmed to be of the r1 class. By generation 8, one of the two had spread to more than 90% frequency yet the dominant allele was reversed in the replicate cages – suggesting that selection for one or the other resistant mutations is stochastic and not because one is more effective at restoring fertility. The equivalent release of *vas2-CRISPR*^*h*^ generated 9-12 mutant alleles above 1% frequency by generation 2 and this variance was maintained over time despite a strong stratification towards those of the r1 class [9]. This reduced complexity of alleles generated by *zpg-CRISPR*^*h*^ may be related to the late onset of resistance, when the majority of alleles at the target locus are drive alleles meaning that there are very few ‘free’ alleles on which r1 alleles can be generated, resulting in the stochastic selection of just a few.

The very late onset of resistance – if it is to occur – is a feature, not immediately intuitive, of this type of suppression drive. Selection for r1 alleles only really becomes strong when the gene drive is close to fixation – when most individuals have at least one copy of the gene drive the relative advantage of an r1 allele over remaining WT alleles (that have a high probability of being removed by homing of the gene drive allele) or the drive allele (with high probability of finding itself in a barren female, homozygous for the drive) is at its highest.

### Incorporating experimental rates of resistance, embryonic end-joining and homology directed repair into a population model

We extend previous models of homing-based gene drives [9, 33, 34], allowing the option of embryonic activity from paternally and maternally derived nuclease that can be resolved through end-joining – forming r1 and r2 alleles – or HDR. Using this model (see **Supplementary Methods**) and the baseline parameter values from rates of fecundity and repair outcomes following germline and embryonic nuclease activity (**Supp Tables 1 and 2**) we generated the expected dynamics of spread and population egg output over the 16 generations of the experiment (**Figure 6**). For both 50% and 10% releases, the model captures the general trend of initial spread that is met by a significant suppression in egg output and an eventual recovery of the population as the gene drive is replaced by resistant mutations. Notably, the observed dynamics of spread and population suppression were faster than the deterministic prediction in all replicates. This may be explained by sensitivity of the model to the fertility of female *zpg-CRISPR*^*h*^ heterozygotes and the extent to which fertility is restored by resistant R1 alleles (**Supplementary Figure 2**) – two parameters that are particularly difficult to measure experimentally [7, 9]. Interestingly, these parameters exert their effect upon drive dynamics in concert: improvements to heterozygous female fertility increase the rate of spread and peak in drive frequency, whereas an r1 allele that fails to restore full wild-type fertility retards the rate at which the drive is removed from the population.

**Figure 6.**
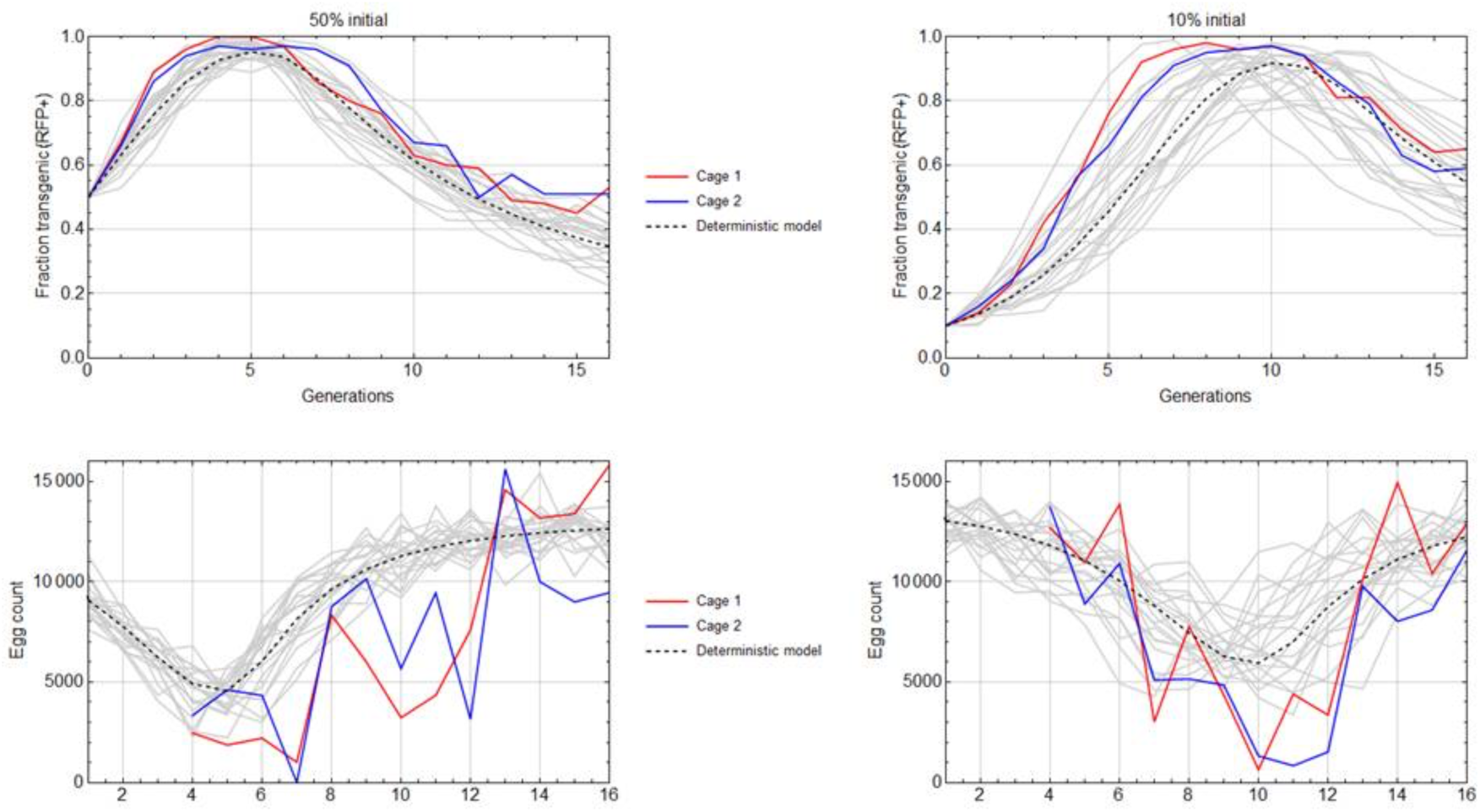
Comparison of observed *zpg-CRISPR*^*h*^ drive frequency and egg output data to deterministic and stochastic model predictions. *zpg-CRISPR*^*h*^/*+* individuals were released into replicate caged WT populations at 50% (left) or 10% (right) initial frequency of *CRISPR*^*h*^/*+*. The frequency of drive-modified mosquitoes (top panels) and the entire egg output per generation (bottom panels) were recorded. Observed data were compared to a deterministic, discrete-generation model (dashed line) based upon observed rates of fertility, homing, and resistance generated by end-joining in the germline and embryo (Supp Table S1). Grey lines show 20 stochastic simulations assuming that females may fail to mate, or mate once with a male of a given genotype according to its frequency in the male population; mated females produce eggs in amounts determined by sampling with replacement from experimental values that subsequently hatch according to the genotype of the mother; of the 600 larvae chosen randomly to populate next generation, some fail to survive to adulthood. Probabilities of mating, egg production, hatching and survival from larval stage to adult are estimated from experiment (Supp Table S2) and random numbers for these events drawn from the appropriate multinomial distributions. Model egg counts for each release are normalised such that the deterministic value corresponds to the average of the two replicate cage experiments at generation sixteen.

## Conclusions

Building suppression gene drives for mosquito control that are fit for purpose requires a combination of the following features: optimal target sites that show high functional constraint; the targeting of multiple sites by the same gene drive construct; fine tuning of the expression of the nuclease that serves as the gene drive’s ‘motor’ in order that the drive shows the most efficient invasion dynamics. A number of these theorised improvements to design have been substantiated: multiplexing through the use of multiple guide RNAs in a single drive construct has been shown to improve robustness of the drive [12, 20, 35]; judicious choice of functionally constrained sites in essential genes has meant that gene drives targeting such sites do not generate resistant alleles that are selected, at least in the laboratory [8, 35]. For the last aspect – optimising expression – we have shown here that simple changes to the promoter controlling nuclease activity can lead to drastic improvements in the speed of invasion due to two principle reasons: 1) less propensity to induce end-joining repair in the germline and 2) improved fecundity of ‘carrier’ females that contain a single copy of the gene drive and transmit it in a super-Mendelian fashion to their offspring.

The combined effect of the improvements conferred by the *zpg* promoter meant that a gene drive targeting a female fertility gene, at a site otherwise extremely prone to generate resistance [9], could still spread close to fixation while imposing a very strong reproductive load before resistant alleles eventually became selected.

The propensity for maternal and paternal deposition of the nuclease under control of the promoters of the drive constructs is an important variable that we have shown can vary greatly. There is also the possibility that both the end-joining/HDR ratio and the magnitude of parental effect might additionally be locus dependent to some extent [12, 29]. Indeed when the *zpg* promoter was used to control expression of a homing based gene drive at the *doublesex* locus we noticed a paternal effect contributing to reduced fecundity that was not observed here [8].

Our data on the relative contributions of end-joining repair and homology-directed repair in the germline and in the early embryo will be particularly important for the modelling of expected performance of gene drive constructs in mosquito populations, not only for suppressive drives but for so-called population replacement drives that also rely on CRISPR-based homing activity [6, 36]. This data, coupled with the improvements to drive performance conferred by promoter choice, should help expedite the building of effective gene drives for mosquito control.

## Methods

### Amplification of promoter and terminator sequences

In determining the length of upstream region to take from each candidate gene to serve as a promoter we took a maximal region of 2kb upstream of the coding sequence, or the entire intergenic distance to the end of the coding sequence of the neighbouring gene, if shorter. The published *Anopheles gambiae* genome sequence provided in Vectorbase [37] was used as a reference to design primers in order to amplify the promoters and terminators of the three *Anopheles gambiae* genes: *AGAP006098* (*nanos*), *AGAP006241* (*zero population growth*) and *AGAP007365* (*exuperantia*). Using the primers provided in Supplementary Table 3 we performed PCRs on 40 ng of genomic material extracted from wild type mosquitoes of the G3 strain using the Wizard Genomic DNA purification kit (Promega). The primers were modified to contain suitable Gibson assembly overhangs (underlined) for subsequent vector assembly, and an *Xho*I restriction site upstream of the start codon. Promoter and terminator fragments were 2092 bp and 601 bp for *nos*, 1074 bp and 1034 bp for *zpg*, and 849 and 1173 bp for *exu*, respectively. The sequences of all regulatory fragments can be found in Supplementary Table 4

### Generation of CRISPR^h^ drive constructs

We modified available template plasmids used previously in Hammond *et al*. (2016) to replace and test alternative promoters and terminators for expressing the Cas9 protein in the germline of the mosquito. p16501, which was used in that study carried a human optimised Cas9 (hCas9) under the control of the *vas2* promoter and terminator, an RFP cassette under the control of the neuronal *3xP3* promoter and a U6:sgRNA cassette targeting the *AGAP007280* gene in *Anopheles gambiae*.

The hCas9 fragment and backbone (sequence containing 3xP3∷RFP and a U6∷gRNA cassette), were excised from plasmid p16501 using the restriction enzymes XhoI+PacI and AscI+AgeI respectively. Gel electrophoresis fragments were then re-assembled with PCR amplified promoter and terminator sequences of *zpg, nos* or *exu* by Gibson assembly to create new *CRISPR*^*h*^ vectors named p17301 *(nos*), p17401 *(zpg)* and p17501 *(exu)*.

### Integration of gene drive constructs at the *AGAP007280* locus

*CRISPR*^*h*^ constructs containing Cas9 under control of the *zpg, nos* and *exu* promoters were inserted into an *hdrGFP* docking site previously generated at the target site in *AGAP007280* (Hammond *et al*., 2016). Briefly, *Anopheles gambiae* mosquitoes of the *hdrGFP-7280* strain were reared under standard conditions of 80% relative humidity and 28°C, and freshly laid embryos used for microinjections as described before[38]. Recombinase-mediated cassette exchange (RCME) reactions were performed by injecting each of the new *CRISPR*^*h*^ constructs into embryos of the *hdrGFP* docking line that was previously generated at the target site in *AGAP007280* [7]. For each construct, embryos were injected with solution containing *CRISPR*^*h*^ (400ng/μl) and a *vas2∷integrase* helper plasmid (400ng/μl)[39]. Surviving G_0_ larvae were crossed to wild type and transformants were identified by a change from GFP (present in the *hdrGFP* docking site) to DsRed linked to the *CRISPR*^*h*^ construct that should indicate successful RCME.

### Phenotypic assays to measure fertility and rates of homing

Heterozygous *CRISPR*^*h*^/+ mosquitoes from each of the three new lines *zpg-CRISPR*^*h*^, *nos-CRISPR*^*h*^, *zpg-CRISPR*^*h*^, were mated to an equal number of wild type mosquitoes for 5 days in reciprocal male and female crosses. Females were blood fed on anesthetized mice on the sixth day and after 3 days, a minimum of 40 were allowed to lay individually into a 25-ml cup filled with water and lined with filter paper. The entire larval progeny of each individual was counted and a minimum of 50 larvae were screened to determine the frequency of the DsRed that is linked to the *CRISPR*^*h*^ allele by using a Nikon inverted fluorescence microscope (Eclipse TE200). Females that failed to give progeny and had no evidence of sperm in their spermathecae were excluded from the analysis. Statistical differences between genotypes were assessed using the Kruskal-Wallis and Mann-Whitney tests.

### Caged Population Invasion Experiments

The cage trials were performed following the same principle described before in Hammond *et al*. (2016). Briefly, heterozygous *zpg-CRISPR*^*h*^ that had inherited the drive from a female parent were mixed with age-matched wild type at L1 at 10% or 50% frequency of heterozygotes. At the pupal stage, 600 were selected to initiate replicate cages for each initial release frequency. Adult mosquitoes were left to mate for 5 days before they were blood fed on anesthetized mice. Two days after, the mosquitoes were left to lay in a 300 ml egg bowl filled with water and lined with filter paper. Each generation, all eggs were allowed two days to hatch and 600 randomly selected larvae were screened to determine the transgenic rate by presence of DsRed and then used to seed the next generation. From generation 4 onwards, adults were blood-fed a second time and the entire egg output photographed and counted using JMicroVision V1.27. Larvae were reared in 2L trays in 500ml of water, allowing a density of 200 larvae per tray. After recovering progeny, the entire adult population was collected and entire samples from generations 0, 2, 4, 5, 6, 7, 8 and 12 (50% release) and 0, 2, 5, 8, 9, 10, 11, 12 and 15 (10% release) were used for pooled amplicon sequence analysis.

### Pooled amplicon sequencing

Pooled amplicon sequencing was performed essentially as described before in Hammond *et al*. (2017). Genomic DNA was mass extracted from pooled samples of mosquitoes using the Wizard Genomic DNA purification kit (Promega), and 90 ng of each used for PCR using KAPA HiFi HotStart Ready Mix PCR kit (Kapa Biosystems) in 50 ul reactions. For caged experiment generations 0, 2, 5 & 8, a 332 bp locus spanning the target site was amplified using primers designed to include the Illumina Nextera Transposase Adapters (underlined), 7280-Illumina-F (<u>TCGTCGGCAGCGTCAGATGTGTATAAGAGACAG</u>GGAGAAGGTAAATGCGCCAC) and 7280-Illumina-R (<u>GTCTCGTGGGCTCGGAGATGTGTATAAGAGACAG</u>GCGCTTCTACACTCGCTTCT). For caged experiment generations 4, 6, 7 and 12 of the 50% release; and 9, 10, 11, 12 and 15 of the 10% release, a 196 bp locus spanning the target site was amplified using primers were designed to include Illumina partial adapters (underlined), Illumina-AmpEZ-7280-F1 (<u>ACACTCTTTCCCTACACGACGCTCTTCCGATCT</u>CGTTAACTGTCTTGGTGGTGAGG) and Illumina-AmpEZ-7280-R1 (<u>GACTGGAGTTCAGACGTGTGCTCTTCCGATCT</u>CACGCTTAACGTCGTCGTTTC). For deposition testing, a 200 bp locus spanning the target site was amplified using primers designed to include Illumina partial adapters (underlined), Illumina-AmpEZ-7280-F2 (<u>ACACTCTTTCCCTACACGACGCTCTTCCGATCT</u>CGGGCAAGAAGTGTAACGG) and Illumina-AmpEZ-7280-R2 (<u>TGGAGTTCAGACGTGTGCTCTTCCGATCTGT</u>CGTTTCTTCCGATGTGAAC). PCR reactions were amplified for 20 cycles and subsequently processed and sequenced using an Illumina MiSeq instrument (Genewiz).

### Analysis of pooled amplicon sequencing

Pooled amplicon sequencing reads were analysed using CRISPResso software v1.0.8 [40] using an average read quality threshold of 30. Insertions and deletions were included if they altered a window of 20 bp surrounding the cleavage site that was chosen on the basis of previously observed mutations at this locus [9]. Allele frequencies were calculated by summing individual insertion or deletion events across all haplotypes on which they were found. A large insertion event, representing incomplete homing of *CRISPR*^*h*^, was found to occur outside of this window and its combined frequency across several haplotypes was calculated and included in the final frequency tables.

### Deposition Testing

F1 heterozygotes containing a gene drive and resistant allele (GD/r1) were generated by crossing *zpg*-*CRISPR*^*h*^, *nos-CRISPR*^*h*^ or *vas2*-*CRISPR*^*h*^ to a resistant strain that is homozygous for the 203-GAGGAG r1 allele at the target site in *AGAP007280*. This scheme allowed the testing of all gene drive constructs, including those (e.g. *vas2*-*CRISPR*^*h*^)would otherwise cause high levels of female sterility when balanced against a wild type allele” An average of 40 F1 heterozygotes were group mated to an excess of wild-type in reciprocal male and female crosses, and allowed to lay *en masse*. F2 progeny were screened for the absence of DsRed that is linked to the *CRISPR*^*h*^ allele and pooled together for mass genomic DNA extraction and pooled amplicon sequencing as described elsewhere. A minimum of 118 F2 individuals were interrogated for each condition.

### Estimation of Meiotic End-joining

A minimum of 200 male or 500 female *zpg*-*CRISPR*^*h*^ drive heterozygotes (F0) were crossed to a line heterozygous for a GFP+ marked knock-out *AGAP007280* allele and allowed to lay eggs. Upon egg-hatching, L1 larvae were sorted using COPAS (complex object parametric analyser and sorter), as in Marois *et al*. (2012). The very rare fraction of progeny (F1) that inherited the GFP+ marked knockout but not the *zpg*-*CRISPR*^*h*^ allele due to lack of homing, were isolated from the progeny pool. They were grown to adulthood and their gDNA was individually extracted using either the Wizard Genomic DNA purification kit (Promega) or the DNeasy Blood & Tissue Kit (Qiagen). A 1048 bp region spanning the gene drive target site was amplified using primers Seq-7280-F (GCACAAATCCGATCGTGACA) and Seq-7280-R3 (GGCTTCCAGTGGCAGTTCCGTA) and Sanger-sequenced using primer Seq-7280-F5 (CGTTTGTGTGTCAGAGCAAGTCG), so that only the allele that was exposed to prior nuclease activity meiotically would amplify (and not the GFP+ allele). In total, 219 F1 progeny descended from female *zpg*-*CRISPR*^*h*^ heterozygote and 240 F1 progeny descended from male *zpg*-*CRISPR*^*h*^ heterozygote were analysed.

### Ethics statement

All animal work was conducted according to UK Home Office Regulations and approved under Home Office License PPL 70/6453.

---

**Supplementary Figure 1.**
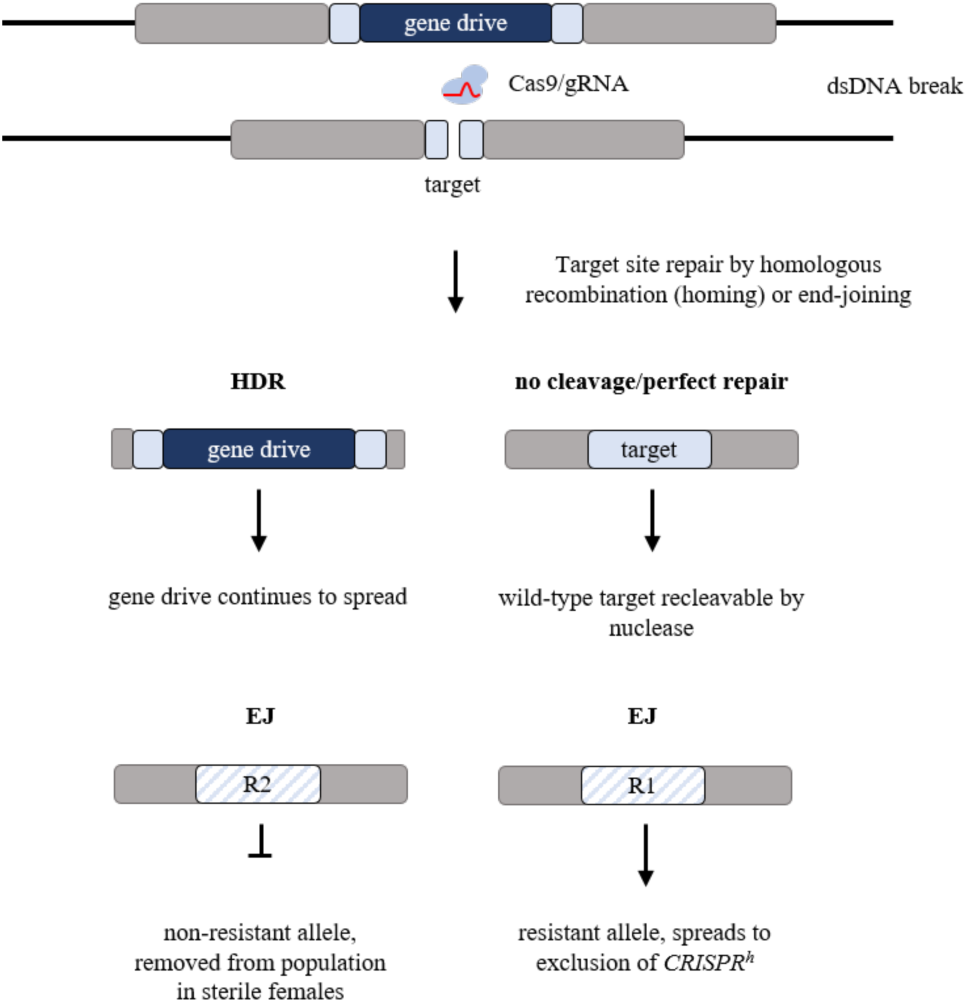
Cleavage by *CRISPR*^*h*^ can generate resistant mutations as a by-product of error-prone end-joining. After cleavage by the nuclease, the majority of target sites will be repaired by homology-directed repair (HDR), leading to a copying over of the *CRISPR*^*h*^ allele called homing. A small fraction of targets may remain unmodified or may repair perfectly, resulting in a target that can be cleaved upon re-exposure by the nuclease. Chromosomes that are repaired by end-joining may generate a mutant target site that can no longer be cleaved by the nuclease. If the target site is essential (i.e. a female fertility gene), then a mutation that disrupts the function of the target gene, called an R2 mutation, will be selected out of the population. Mutations that re-code a functional target gene, called an R1 mutation, are resistant to the gene drive and will come under strong selection in presence of the drive.

**Supplementary Figure 2.**
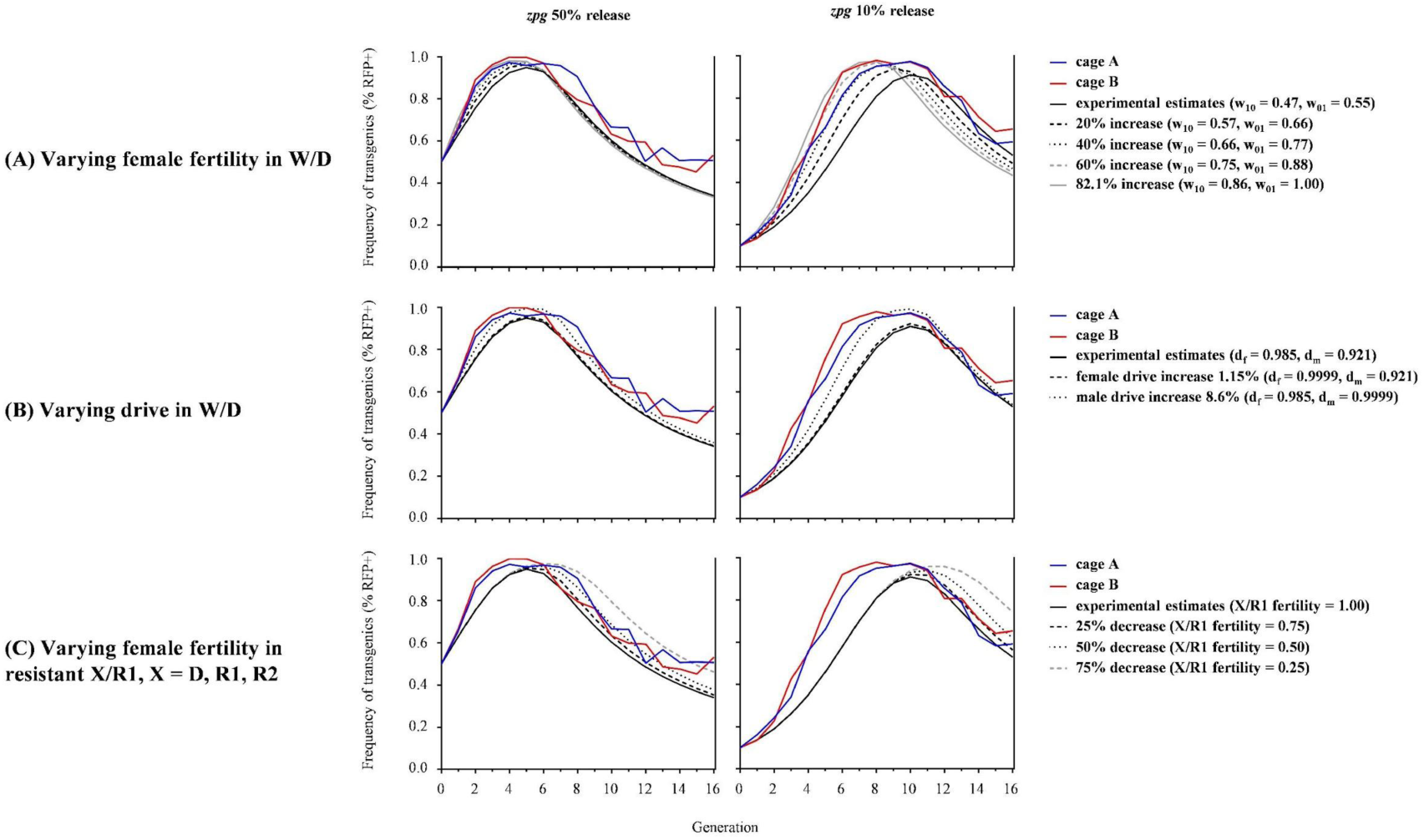
Comparison of cage data with deterministic model results incorporating percentage change in parameter estimates. Note that for (A), W/D female fertilities with maternal/paternal parental effects are varied together, keeping their ratio constant.

**Supplementary Figure 3.**
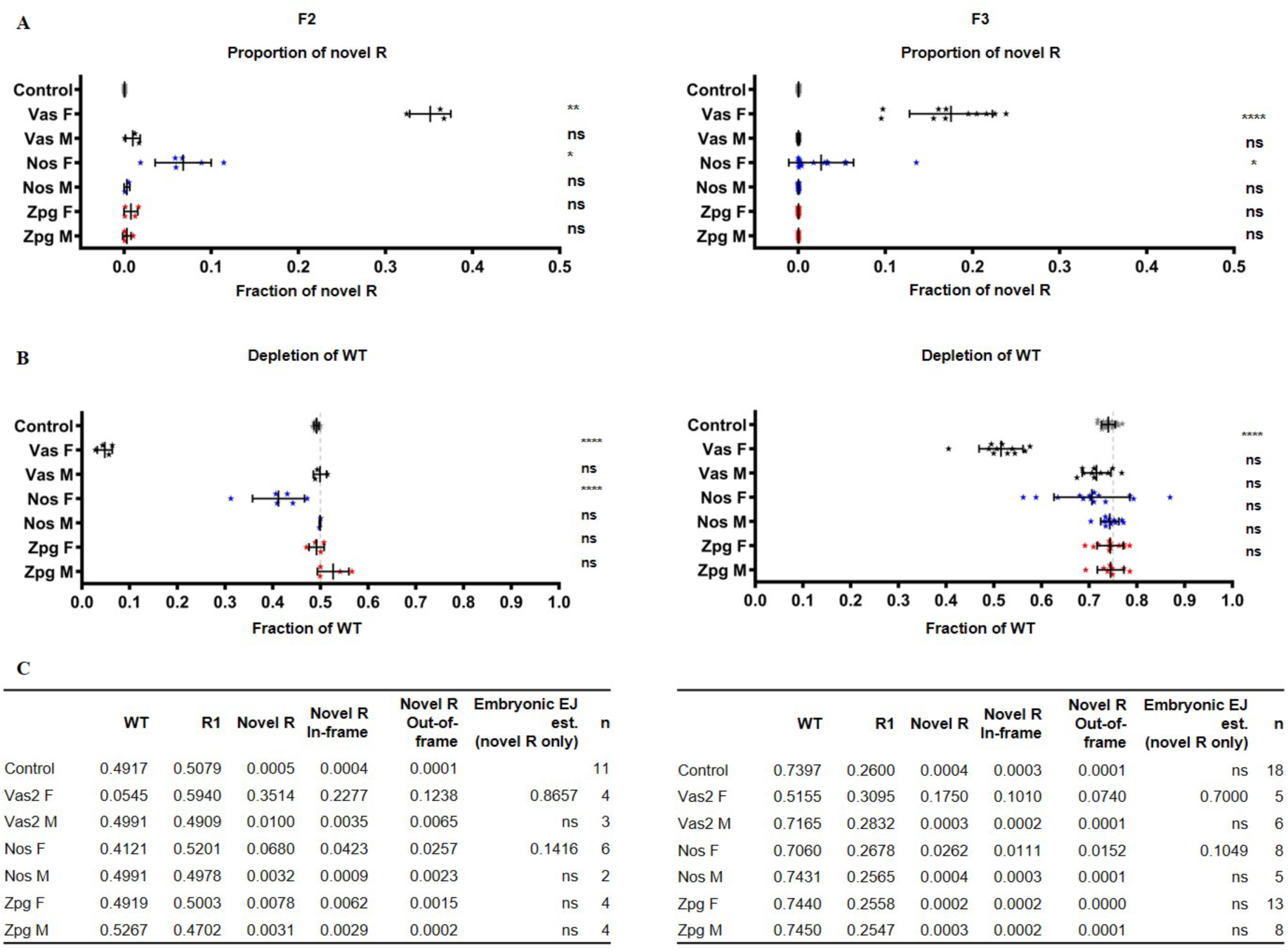
Depletion of wild-type and formation of mutations caused by parentally deposited nuclease from different gene drive constructs. *zpg-CRISPR*^*h*^, *nos-CRISPR*^*h*^ or *vas2-CRISPR*^*h*^ were crossed to a homozygous resistant strain (r1 203-GAGGAG) to generate F1 heterozygotes containing both a gene drive and resistant allele (GD/r1). F1 heterozygotes and r1/r1 homozygotes (control) were crossed to wild type and their non-drive F2 progeny analysed by pooled amplicon sequencing across the target site in AGAP007280. Amplicon sequencing results from F2 individuals (A) with genotype wt/R1 (left) and their F3 progeny (B) deriving from the F2 crossed to wild type (right) reveal novel R alleles (top), depletion of wt (middle). Frequencies expected by Medelian inheritance of the wt allele are indicated (dashed line). Deposited nuclease significantly depleted wt and generated novel R for *nos-CRISPR*^*h*^ and *vas2-CRISPR*^*h*^ females in the F2, but not *zpg-CRISPR*^*h*^ females, or males of any class (Brown-Forsythe and Welch ANOVA, ****: *p* < 0.0001, ***: *p* < 0.001, *: *p* < 0.05, ns: non-significant). Rates of embryonic EJ (tables) were estaminated by calculating the frequency of novel R amongst WT and novel R (in the F2), or by multiplying the frequency of novel R by 4 (in the F3), the dilution factor expected due to additional wt alleles received from the parents.

## Supplementary Methods and Tables

### Supplementary Methods

#### Population genetics model

To model the results of the cage experiments, we use discrete-generation recursion equations for the genotype frequencies, treating males and females separately. *F*_*i*_(*t*) and *M*_*i*_(*t*) denote the frequency of females (or males) of genotype *i* = *X*/*Y* in the total female (or male) population. We consider four alleles, W (wildtype), D (driver), R_1_ (functional resistant) and R_2_ (non-functional resistant), and therefore ten basic genotypes. Both resistant alleles cause a change in the target sequence such that it is no longer recognised by the nuclease, but function of the target gene is restored fully or partially in R_1_ alleles and destroyed in R_2_ alleles.

##### Parental effects

We consider that further cleavage of the W allele and repair can occur in the embryo if nuclease is present, due to one or both contributing gametes derived from a parent with one or two driver alleles. Previously, embryonic end-joining was modelled as acting immediately in the zygote [1,2]; we consider here that individuals may be mosaics with intermediate phenotypes and therefore we model embryonic activity as causing mosaicism in both the soma, affecting female fitness [3], and also in germline cells, altering gene transmission [4]. No correlation is assumed between the effect on fitness and the effect on gene transmission.

Extending our models [3,4] to include two types of resistance (R_1_and R_2_), we denote mosaic individuals with parental effects (i.e., on fitness and gene transmission rates) as W/W(*a*), W/D(*a*), W/R_1_(*a*) and W/R_2_(*a*), where *a* = 10, 01 or 11 denotes nuclease from the mother, father or both. Individuals without parental effects are denoted only as X/Y. Gametes are distinguished by whether they carry deposited parental nuclease: W, R_1_, and R_2_ gametes are derived from parents that have no drive allele and therefore carry no deposited nuclease, and gametes 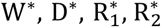 are derived from individuals with one or two copies of the drive allele and carry nuclease that is transmitted to the zygote. For example, W/R_1_(10) mosaic individuals start as zygotes that have either a W* or R_1_* egg from the mother and correspondingly R_1_ or W sperm from the father, and due to the deposited nuclease, the wildtype allele may undergo further cleavage and embryonic end-joining and HDR in the embryo.

##### Fitness

Let *w*_*i*_ ≤ 1 represent the fitness of genotype *i* = X/Y relative to *w*_ww_ = 1 for the wild-type homozygote. We assume no fitness effects in males. Fitness effects in females are manifested as differences in the relative ability of genotypes to participate in reproduction. We assume the target gene is needed for female participation in reproduction, thus D/D, D/R_2_, and R_2_/R_2_ females do not reproduce, and there is no reduction in fertility in females with only one copy of the gene if no parental effects are present. To model parental effects on fitness (as in [3,4]), genotypes with parental nuclease W/W(*a*), W/D(*a*) and W/R_2_(*a*), *a* = 10, 01 or 11, are assigned an intermediate fitness 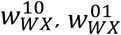, or 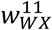 depending on whether nuclease was derived from a transgenic mother, father, or both (no reduction in fitness of W/R_1_ females with parental nuclease, because all cells in the soma have at least one functioning copy of the target gene). We assume that parental effects are the same whether the parent(s) had one or two drive alleles. For simplicity, the same baseline reduced fitness of *w*_11_, *w*_01_, *w*_11_ is assigned to all mosaic genotypes W/W(*a*), W/D_1_(*a*) and W/R_2_(*a*) with maternal, paternal and maternal/paternal effects, *a* = 10, 01 or 11, with fitness estimated as the product of mean egg production values and hatching rates relative to wild-type in Supp. Table 1 (deterministic model).

##### Gene transmission

We model parental effects on rates of gene transmission from mosaics W/W(*a*), W/D(*a*), W/R_1_(*a*) and W/R_2_(*a*), *a* = 10, 01 or 11, by assuming that parentally-derived nuclease can be active in the germline, leading to mosaicism that affects the types and proportions of gametes contributed.

Overall gene transmission from drive individuals includes both greater-than-Mendelian inheritance of the drive allele from germline W/D cells and possible parental effects due to nuclease deposition in the embryo. To incorporate observed measurements of drive into the model, we define overall rates of transmission from W/D individuals, with the proportions of gametes 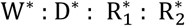 (all carry parental nuclease) from W/D(*a*) individuals depending upon whether the deposited nuclease is from the mother, father, or both (*a* = 10, 01 or 11):

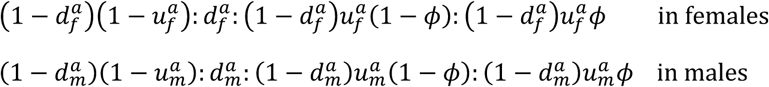

Here, 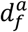 and 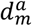 are the observed rates of transmission of the driver allele in the two sexes, 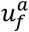 and 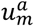 are the fractions of non-drive gametes that are resistant (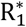 and 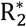 alleles), and *ϕ* is the fraction of resistant alleles that are non-functional 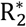 alleles.

In mosaic types W/W(*a*), W/R_1_(*a*) and W/R_2_(*a*) with *a* = 10, 01 or 11 for parental effects due to nuclease deposition from mother, father or both, we define the proportion of wild-type alleles in the germline stem cells that are cleaved and repaired to resistant alleles (R_1_ and R_2_) by end joining as *δ*_10_, *δ*_01_, *δ*_11_ with nuclease from mother, father or both, and the proportion repaired by HDR as *ε*_10_, *ε*_01_, *ε*_11_. These parameters are estimated from deposition experiments on W/R1 individuals. Supp. Table S5 shows the resulting proportions of different genotypes among germline stem cells (rows) in gonads of mosaic types W/W(*a*), W/R_1_(*a*) and W/R_2_(*a*), *a* = 10, 01 or 11 (columns). Gamete production from W/W, W/R_1_ and W/R_2_ germline stem cells in mosaic individuals is assumed to be Mendelian, since we assume that parental nuclease is no longer active. The resulting proportions of gametes contributed from each type of individual is summarized in Supp. Table 6, along with the fitness for each.

##### Recursion equations

We now consider the gamete contributions from each genotype, including parental effects on fitness and gene transmission. The proportion *e*_*k*_(*t*) (and *s*_*k*_(*t*)) of type *k* gametes in eggs (and sperm) produced by females (and males) participating in reproduction is given by:

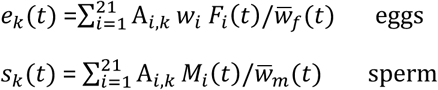

Above, *F*_*i*_(*t*) and *M*_*i*_(*t*) correspond to the frequencies of female/male individuals of type *i* in the female/male population, where *i* is summed over the twenty-one individual types (this includes genotypes *F*_*XY*_(*t*) and *M*_*XY*_(*t*) without parental effects: W/W, W/R_1_, W/ R_2_, D/D, D/R_1_, D/ R_2_, R_1_/R_1_, R_1_/R_2_, and R_2_/R_2_ and also mosaics 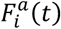 and 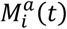 of genotypes *i* = W/W, W/D, W/R_1_ and W/R_2_ distinguished according to parental effect, *a* = 10, 01 or 11). The seven gamete types are distinguished by both the allele that they carry and whether they carry deposited nuclease: *k* = W, R_1_, and R_2_ (without parental nuclease) and 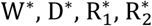 (with parental nuclease). The coefficients *A*_*i,k*_ correspond to the proportion of gametes of type *k* from individuals of type *i* and are given in Supp. Table S6, with row *i* corresponding to an individual of type *i* and columns to proportions of gametes of type *k*. Above, *w*_*i*_ is the fitness of individual of type *i*, where we have assumed all male fitnesses are one, and 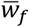 and 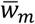 are the average female and male fitnesses:

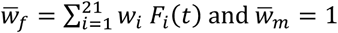

To model cage experiments, we start with an equal number of males and females and an initial starting frequency of heterozygote drive females and males that inherited the drive from their mothers of: 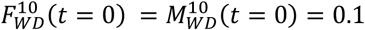 (for 10% release) or 0.5 (for 50% release). The remaining starting population is wildtype only. Assuming a 50:50 ratio of males and females in progeny, after the starting generation, genotype frequencies of type *i* in the next generation (*t* + 1) are the same in males and females, *F*_*i*_(*t* + 1) = *M*_*i*_(*t* + 1). Both are both given by *G*_*i*_(*t* + 1) in the following set of equations in terms of the gamete proportions in the previous generation, assuming random mating:

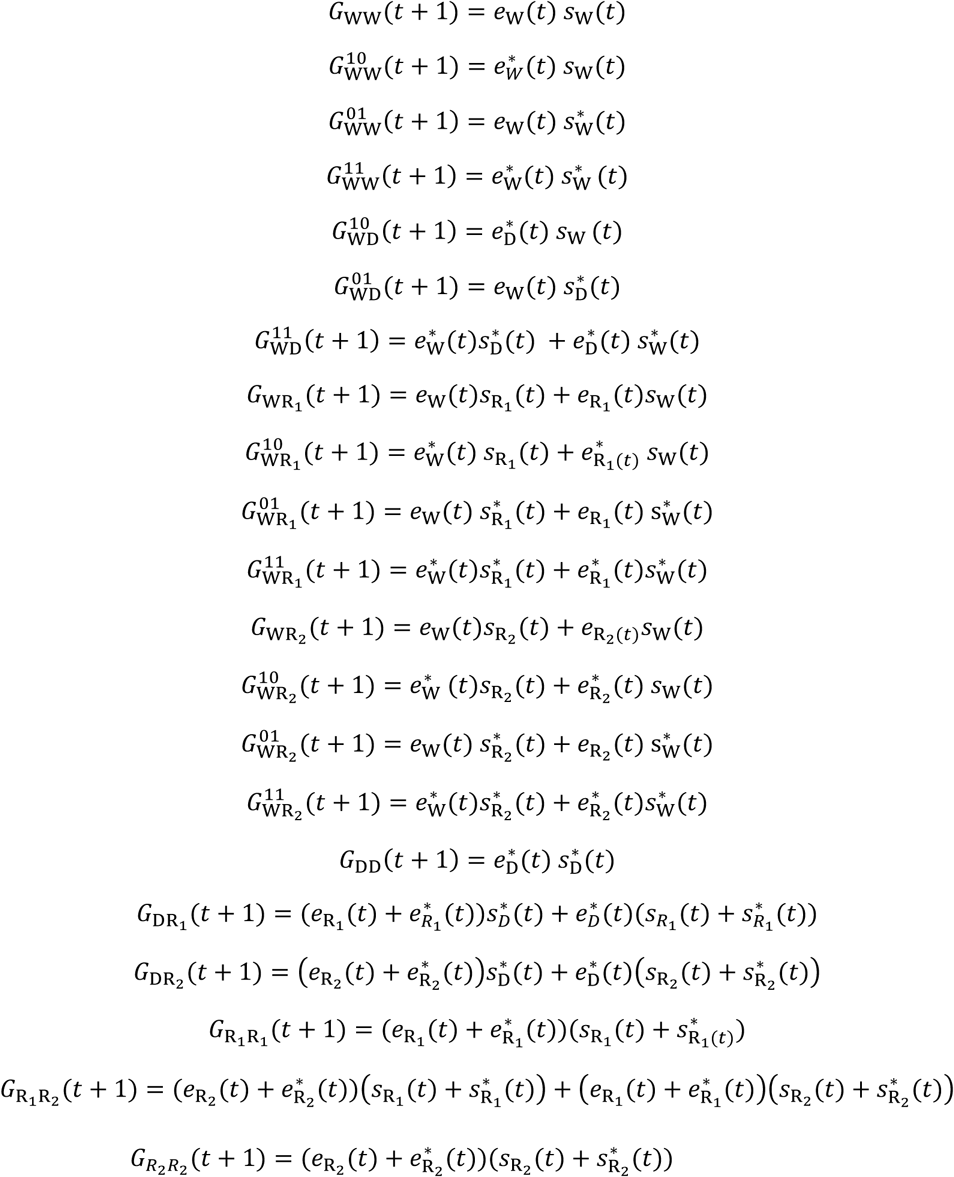

The frequency of transgenic individuals can be compared with experiment (fraction of RFP+ individuals), where here (*t*) is omitted for brevity:

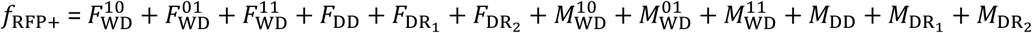

##### Stochastic version

In the stochastic version of the model described above, random values for probabilistic events are taken from the appropriate multinomial distributions, with probabilities estimated from experiment where applicable (Supp. Table 2). To model the cage experiments, 150 female and 150 male wildtype adults (or 270 females and 270 males for 10% release) along with 150 female and 150 male heterozygotes (or 30 females and 30 males for 10% release) are initially present. Females may fail to mate, or mate once in their life, with a male of a given genotype according to its frequency in the male population, chosen randomly with replacement such that males may mate multiple times. The number of eggs produced from each mated female is randomly chosen by sampling with replacement from experimental values, and the eggs hatch or not with a probability that depends on the mother (Supp. Table 2). To start the next generation, 600 larvae are randomly selected, unless less than 600 larvae have hatched, in which case the smaller amount initiates the next generation, following experiment. The probability of subsequent survival to adulthood is assumed to be equal across genotypes. Assuming very large population sizes gives results for the genotype frequencies that are indistinguishable from the deterministic model. For the deterministic egg count, we use the large population limit of the stochastic model.

All calculations are carried out using Wolfram Mathematica [5].

---

**Supp Table 1.** Model parameters for zpg-CRISPR^h^. For combined maternal and paternal effects (nuclease from both parents), the minimum of observed values for maternal or paternal effect is used for the fitness (*w*_11_) and drive (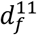 and 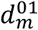); for embryonic HDR and EJ parameters *ϵ*_11_ and *δ* _11_, we use the maximum observed value of each from maternal or paternal effect. We assume that parental effects on fitness and embryonic HDR and EJ parameters for non-drive (W/W, W/R_2_) females with nuclease from one or both parents are the same as observed values for drive heterozygote (W/D) females with parental effects.

| Parameter | Symbol | Estimate | Method of estimation |
| --- | --- | --- | --- |
| Average fertility<br>W/D females from female | $w_{10}$ | 0.471<sup>‡</sup> | Phenotype assay (mean larvae count, relative to WT value) |
| Average fertility<br>W/D females from male | $w_{01}$ | 0.549<sup>‡</sup> | Phenotype assay (mean larvae count, relative to WT value) |
| Drive in<br>W/D females from female | $d_f^{10}$ | 0.985 | Phenotype assay |
| Drive in<br>W/D females from male | $d_f^{01}$ | 0.985 | Phenotype assay |
| Drive in<br>W/D males from female | $d_m^{10}$ | 0.921 | Phenotype assay |
| Drive in<br>W/D males from male | $d_m^{01}$ | 0.921 | Phenotype assay |
| EJ parameter in W/D (fraction non-drive alleles that are resistant) | $u^*$ | 0.392 | Sequencing non-drive progeny of zpg-CRISPR<sup>h</sup> heterozygotes |
| Fraction resistant alleles that are non-functional | $\varphi$ | 0.378 | Single generation resistance assay using zpg-CRISPR<sup>h</sup> |
| Embryonic HDR in<br>W/R1 from female | $\epsilon_{10}$ | 0.0 | Deposition induced EJ and HDR |
| Embryonic HDR in<br>W/R1 from male | $\epsilon_{01}$ | 0.0 | Deposition induced EJ and HDR |
| Embryonic EJ in<br>W/R1 from female | $\delta_{10}$ | 0.0 | Deposition induced EJ and HDR |
| Embryonic EJ in<br>W/R1 from male | $\delta_{01}$ | 0.0 | Deposition induced EJ and HDR |
| Initial frequency of heterozygous drive individuals (that inherited drive from a female parent) | $F_{WD}^{10}, M_{WD}^{10}$ | 0.1 (10% release)<br>0.5 (50% release) | Following cage experiment |

<sup>‡</sup> Deterministic model
\*For comparison to experiment, it is assumed that this parameter is the same in females and males and with maternal or paternal effect, $u_F^{10} = u_F^{01} = u_M^{10} = u_M^{01} = u_M^{11} = u_M^{11} = u$ .

**Supp Table 2.** Additional parameters for stochastic model for zpg-CRISPR^h^. We assume that parental effects on fitness (egg production and hatching rates) for non-drive (W/W, W/R_2_) females with nuclease from one or both parents are the same as observed values for female drive heterozygote (W/D) females with parental effects.

| Parameter | Estimate | Method of estimation |
| --- | --- | --- |
| Mating probability | 0.85 | Hammond et al. 2017 |
| Egg count<br>wildtype | {0, 0, 61, 63, 66, 77, 77, 92, 93, 99, 107, 108, 109, 109, 111, 113, 115, 116, 116, 118, 119, 120, 120, 121, 121, 124, 126, 127, 127, 127, 128, 129, 130, 130, 132, 134, 135, 136, 140, 144, 151, 153, 161, 162, 171} (Mean 113.7) | Phenotype assay |
| Egg count<br>W/D female from female | {0, 0, 0, 0, 17, 17, 20, 20, 39, 42, 44, 45, 47, 47, 50, 59, 60, 60, 61, 61, 67, 67, 77, 80, 88, 88, 94, 97, 104, 107, 110, 111, 117, 119, 122, 148, 154, 155} (Mean 68.3) | Phenotype assay |
| Egg count<br>W/D female from male | {0, 12, 36, 41, 42, 43, 51, 57, 61, 61, 64, 66, 67, 68, 70, 71, 76, 81, 83, 87, 88, 89, 91, 93, 93, 94, 102, 105, 106, 110, 115, 115, 116, 127, 128, 135, 182} (Mean 81.8) | Phenotype assay |
| Hatching rate,<br>wildtype<br>(no parental nuclease) | 0.954 | Phenotype assay (mean larvae count divided by mean egg count) |
| Hatching rate,<br>W/D female from female | 0.748 | Phenotype assay (mean larvae count divided by mean egg count) |
| Hatching rate,<br>W/D female from male | 0.729 | Phenotype assay (mean larvae count divided by mean egg count) |
| Probability of emergence<br>from pupa (survival from larva) | 0.73 | Average over all gens. and cage experiments |
| Initial population (zero generation) | 50% (10%) release: 150 (270) female and 150(270) male WT adults, 150 (30) female and 150 (30) male heterozygous zpg-CRISPR<sup>h</sup> that inherited drive from a female parent. | Following cage experiment |

**Supp Table 3.**
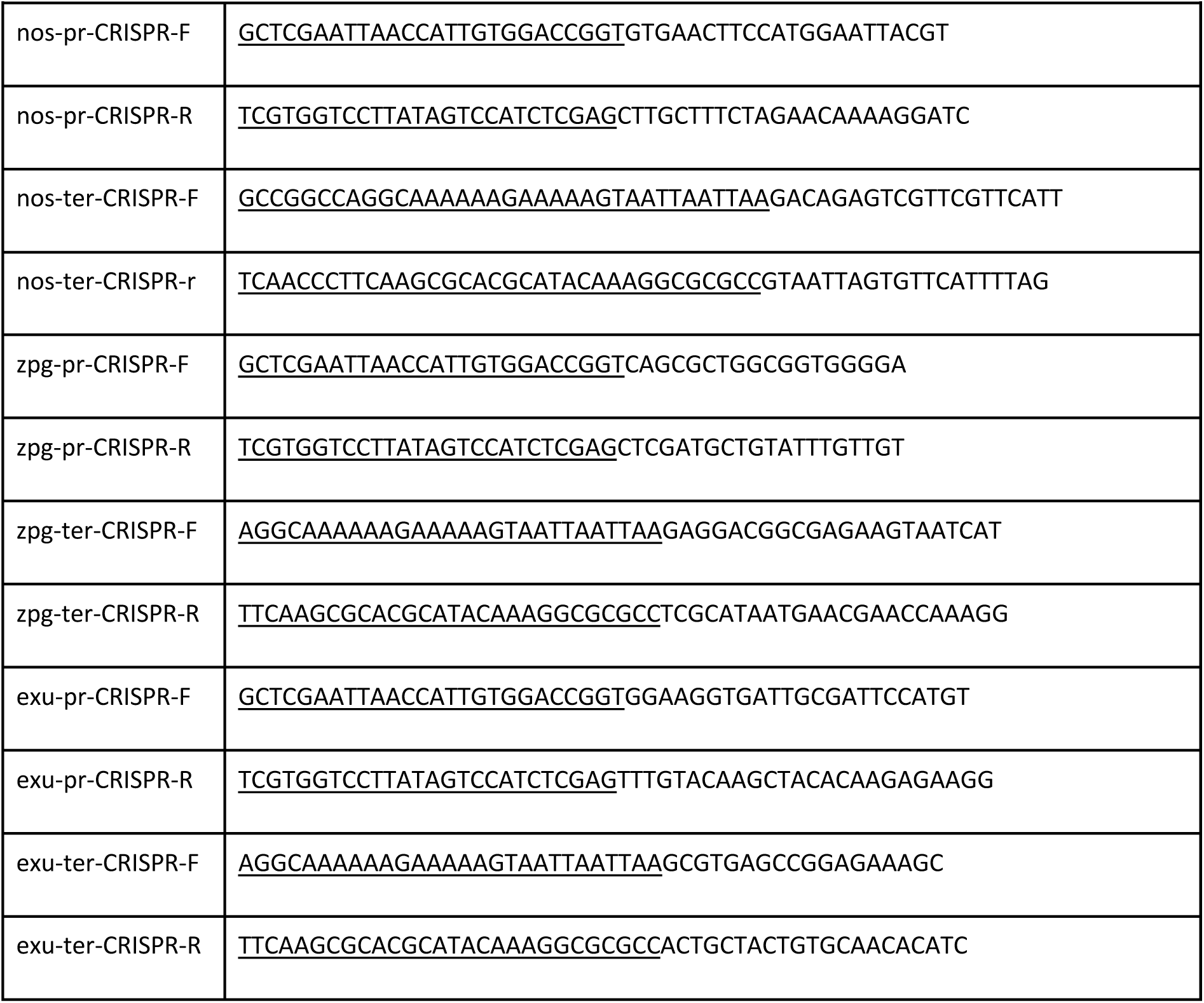
Primers used in this study to assemble the vectors. Primers used to amplify *nos, zpg* and *exu* promoter (pr) and terminator (ter) sequences. Underlined are the Gibson adaptors used to clone promoter and terminator fragments into the CRISPR^h^ vector.

**Supplementary Table 4.**
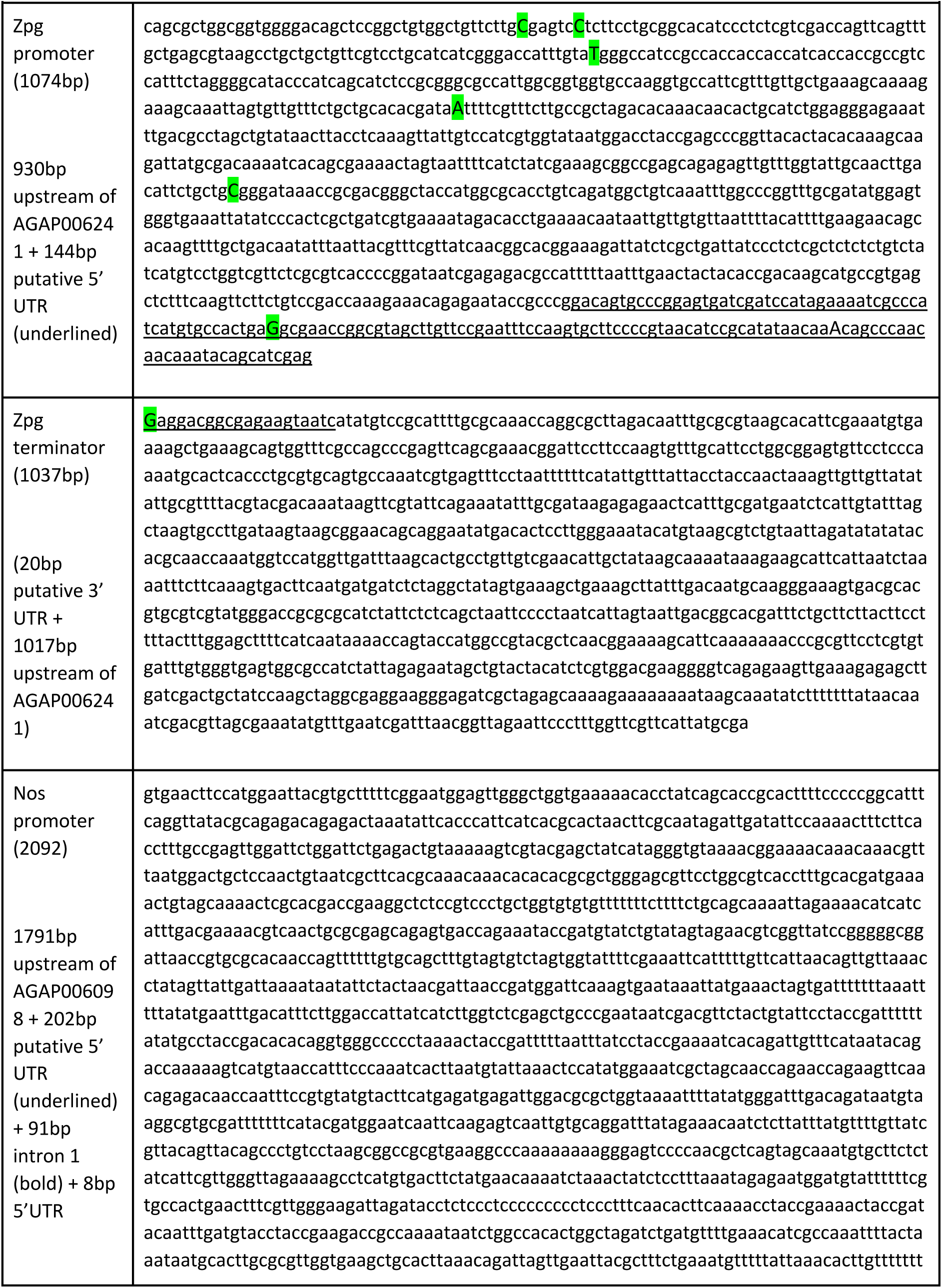
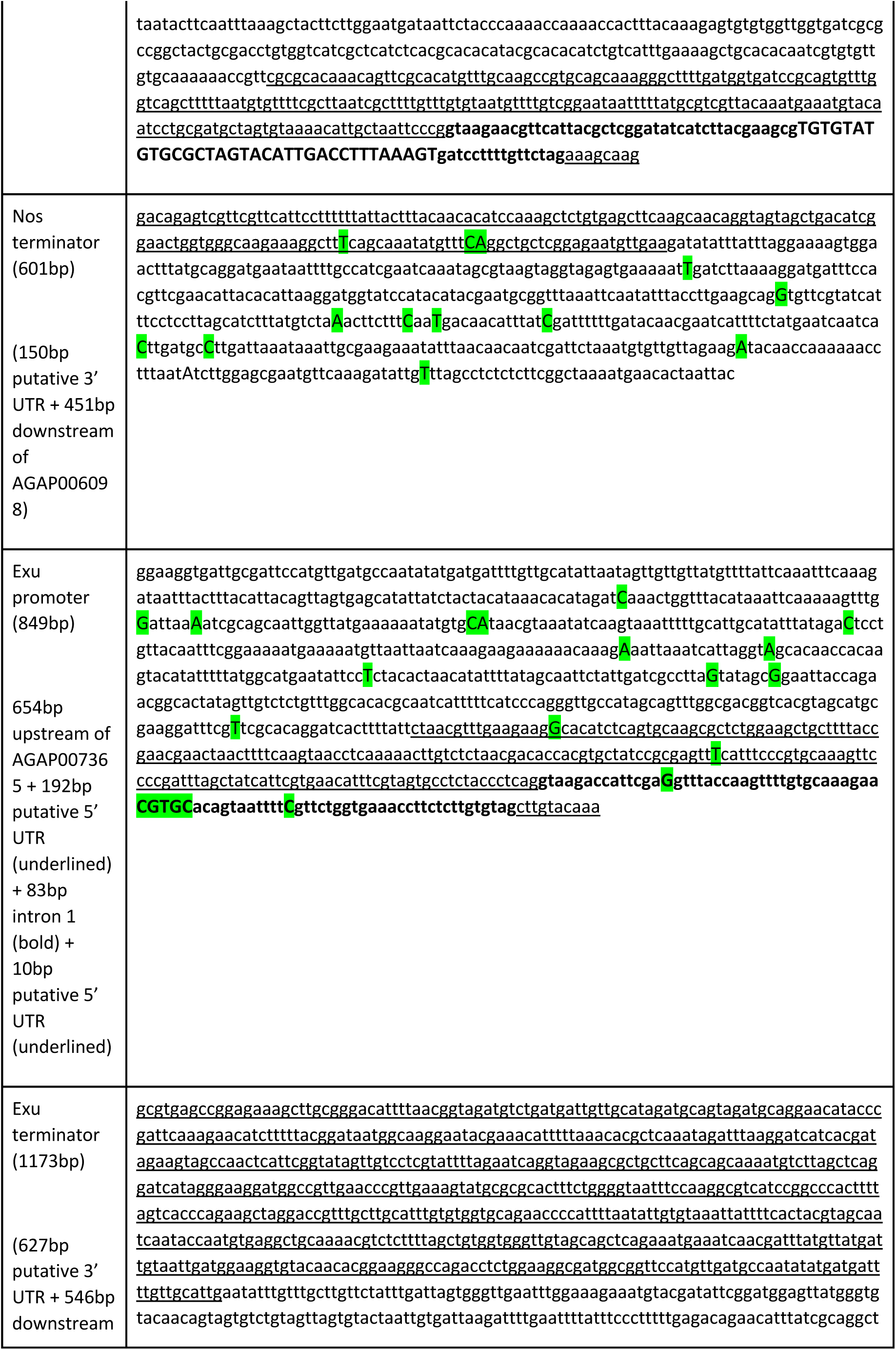
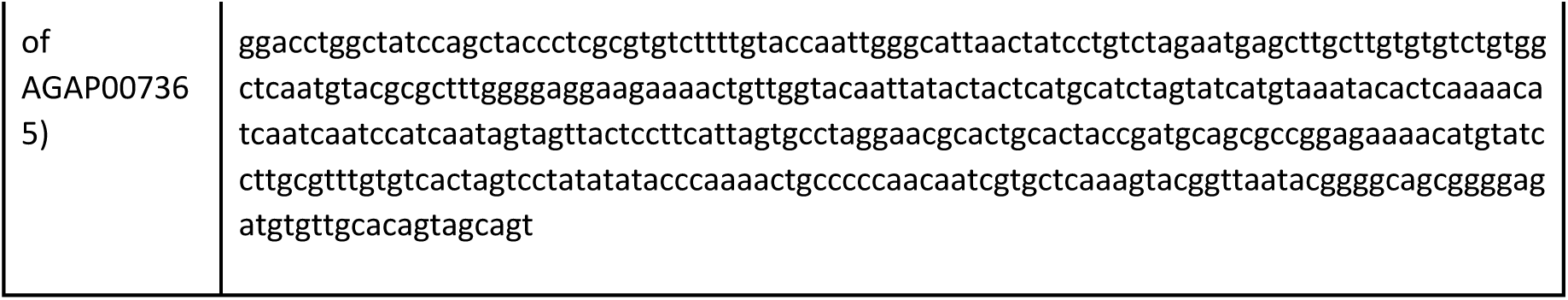
Promoter and terminator sequences.

**Supp Table 5.** Proportions of germline cells in W/Y (Y = W, R_1_, R_2_) with parental effects. For the model of parental effects, the proportions of germline stem cells of different genotypes in female or male (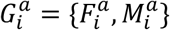 mosaics (derived from zygotes of W/W, W/R_1_ and W/R_2_) where the nuclease is deposited from a mother, father or both (*a* = {10, 01, 11}).

| Germline<br>stem cell | Mosaic genotype |  |  |
| --- | --- | --- | --- |
| | $G_{WW}^a$ | $G_{WR_1}^a$ | $G_{WR_2}^a$ |
| $W/W$ | $(1 - \delta_G^a)^2$ | 0 | 0 |
| $W/R_1$ | $2(1 - \phi_e)(1 - \delta_G^a)\delta_G^a$ | $1 - \varepsilon_G^a - \delta_G^a$ | 0 |
| $W/R_2$ | $2\phi_e(1 - \delta_G^a)\delta_G^a$ | 0 | $1 - \varepsilon_G^a - \delta_G^a$ |
| $R_1/R_1$ | $(1 - \phi_e)^2(\delta_G^a)^2$ | $\varepsilon_G^a + (1 - \phi_e)\delta_G^a$ | 0 |
| $R_1/R_2$ | $2(1 - \phi_e)\phi_e(\delta_G^a)^2$ | $\phi_e\delta_G^a$ | $(1 - \phi_e)\delta_G^a$ |
| $R_2/R_2$ | $(\phi_e)^2(\delta_G^a)^2$ | 0 | $\varepsilon_G^a + \phi_e\delta_G^a$ |

**Supp Table 6.** Genotype fitnesses and the proportion of each type of gamete produced by them. The fitnesses of the 21 types (10 basic genotypes differentiated according to parental effect) and the proportion of each type of gamete produced by them. *G*={*F, M*}, where *F* denotes females and eggs produced; *M* denotes males and sperm produced.

| Genotype ‡ | Fitness | Gametes produced |  |  |  |  |  |  |
| --- | --- | --- | --- | --- | --- | --- | --- | --- |
|  |  | W | R<sub>1</sub> | R<sub>2</sub> | W* | D* | R<sub>1</sub>* | R<sub>2</sub>* |
| W/W | 1 | 1 | 0 | 0 | 0 | 0 | 0 | 0 |
| W/W (10) | $w_{10}$ | $1 - \delta_G^{10}$ | $\delta_G^{10} (1 - \phi_e)$ | $\delta_G^{10} \phi_e$ | 0 | 0 | 0 | 0 |
| W/W (01) | $w_{01}$ | $1 - \delta_G^{01}$ | $\delta_G^{01} (1 - \phi_e)$ | $\delta_G^{01} \phi_e$ | 0 | 0 | 0 | 0 |
| W/W (11) | $w_{11}$ | $1 - \delta_G^{11}$ | $\delta_G^{11} (1 - \phi_e)$ | $\delta_G^{11} \phi_e$ | 0 | 0 | 0 | 0 |
| W/D (10) | $w_{10}$ | 0 | 0 | 0 | $(1 - d_G^{10})(1 - u_G^{10})$ | $d_G^{10}$ | $(1 - d_G^{10})u_G^{10}(1 - \phi)$ | $(1 - d_G^{10})u_G^{10}\phi$ |
| W/D (01) | $w_{01}$ | 0 | 0 | 0 | $(1 - d_G^{01})(1 - u_G^{01})$ | $d_G^{10}$ | $(1 - d_G^{01})u_G^{01}(1 - \phi)$ | $(1 - d_G^{01})u_G^{01}\phi$ |
| W/D (11) | $w_{11}$ | 0 | 0 | 0 | $(1 - d_G^{11})(1 - u_G^{11})$ | $d_G^{10}$ | $(1 - d_G^{11})u_G^{11}(1 - \phi)$ | $(1 - d_G^{11})u_G^{11}\phi$ |
| W/R<sub>1</sub> | 1 | 1/2 | 1/2 | 0 | 0 | 0 | 0 | 0 |
| W/ R<sub>1</sub> (10) | 1 | $(1 - \epsilon_G^{10} - \delta_G^{10})/2$ | $(1 + \epsilon_G^{10} + \delta_G^{10}(1 - \phi_e))/2$ | $\delta_G^{10} \phi_e/2$ | 0 | 0 | 0 | 0 |
| W/ R<sub>1</sub> (01) | 1 | $(1 - \epsilon_G^{01} - \delta_G^{01})/2$ | $(1 + \epsilon_G^{01} + \delta_G^{01}(1 - \phi_e))/2$ | $\delta_G^{01} \phi_e/2$ | 0 | 0 | 0 | 0 |
| W/ R<sub>1</sub> (11) | 1 | $(1 - \epsilon_G^{11} - \delta_G^{11})/2$ | $(1 + \epsilon_G^{11} + \delta_G^{11}(1 - \phi_e))/2$ | $\delta_G^{11} \phi_e/2$ | 0 | 0 | 0 | 0 |
| W/R<sub>2</sub> | 1 | 1/2 | 0 | 1/2 | 0 | 0 | 0 | 0 |
| W/R<sub>2</sub> (10) | $w_{10}$ | $(1 - \epsilon_G^{10} - \delta_G^{10})/2$ | $\delta_G^{10} (1 - \phi_e)/2$ | $(1 + \epsilon_G^{10} + \delta_G^{10} \phi_e)/2$ | 0 | 0 | 0 | 0 |
| W/R<sub>2</sub> (01) | $w_{01}$ | $(1 - \epsilon_G^{01} - \delta_G^{01})/2$ | $\delta_G^{01} (1 - \phi_e)/2$ | $(1 + \epsilon_G^{01} + \delta_G^{01} \phi_e)/2$ | 0 | 0 | 0 | 0 |
| W/R<sub>2</sub> (11) | $w_{11}$ | $(1 - \epsilon_G^{11} - \delta_G^{11})/2$ | $\delta_G^{11} (1 - \phi_e)/2$ | $(1 + \epsilon_G^{11} + \delta_G^{11} \phi_e)/2$ | 0 | 0 | 0 | 0 |
| D/D | 0 | 0 | 0 | 0 | 0 | 1 | 0 | 0 |
| D/ R<sub>1</sub> | 1 | 0 | 0 | 0 | 0 | 1/2 | 1/2 | 0 |
| D/ R<sub>2</sub> | 0 | 0 | 0 | 0 | 0 | 1/2 | 0 | 1/2 |
| R<sub>1</sub>/R<sub>1</sub> | 1 | 0 | 1 | 0 | 0 | 0 | 0 | 0 |
| R<sub>1</sub>/R<sub>2</sub> | 1 | 0 | 1/2 | 1/2 | 0 | 0 | 0 | 0 |
| R<sub>2</sub>/R<sub>2</sub> | 0 | 0 | 0 | 1 | 0 | 0 | 0 | 0 |

$\ddagger(10), (01)$ , or $(11)$ denotes a mosaic type with effects of deposited parental nuclease from mother, father or both.

## References

1. Bhatt S., Weiss D.J., Cameron E., Bisanzio D., Mappin B., Dalrymple U., Battle K., Moyes C.L., Henry A., Eckhoff P.A., et al. 2015 The effect of malaria control on Plasmodium falciparum in Africa between 2000 and 2015. Nature 526(7572), 207–211. (doi: 10.1038/nature15535).

2. World Health Organisation. 2017 Vector-borne diseases. (Geneva.

3. Burt A. 2003 Site-specific selfish genes as tools for the control and genetic engineering of natural populations. Proc Biol Sci 270(1518), 921–928. (doi: 10.1098/rspb.2002.2319).

4. Esvelt K.M., Smidler A.L., Catteruccia F., Church G.M. 2014 Concerning RNA-guided gene drives for the alteration of wild populations. eLife 3, e03401. (doi: 10.7554/eLife.03401).

5. Gantz V.M., Bier E. 2015 Genome editing. The mutagenic chain reaction: a method for converting heterozygous to homozygous mutations. Science 348(6233), 442–444. (doi: 10.1126/science.aaa5945).

6. Gantz V.M., Jasinskiene N., Tatarenkova O., Fazekas A., Macias V.M., Bier E., James A.A. 2015 Highly efficient Cas9-mediated gene drive for population modification of the malaria vector mosquito Anopheles stephensi. Proc Natl Acad Sci U S A 112(49), E6736–6743. (doi: 10.1073/pnas.1521077112).

7. Hammond A., Galizi R., Kyrou K., Simoni A., Siniscalchi C., Katsanos D., Gribble M., Baker D., Marois E., Russell S., et al. 2016 A CRISPR-Cas9 gene drive system targeting female reproduction in the malaria mosquito vector Anopheles gambiae. Nat Biotechnol 34(1), 78–83. (doi: 10.1038/nbt.3439).

8. Kyrou K., Hammond A.M., Galizi R., Kranjc N., Burt A., Beaghton A.K., Nolan T., Crisanti A. 2018 A CRISPR-Cas9 gene drive targeting doublesex causes complete population suppression in caged Anopheles gambiae mosquitoes. Nat Biotechnol 36(11), 1062–1066. (doi: 10.1038/nbt.4245).

9. Hammond A.M., Kyrou K., Bruttini M., North A., Galizi R., Karlsson X., Kranjc N., Carpi F.M., D’Aurizio R., Crisanti A., et al. 2017 The creation and selection of mutations resistant to a gene drive over multiple generations in the malaria mosquito. PLoS Genet 13(10), e1007039. (doi: 10.1371/journal.pgen.1007039).

10. Champer J., Reeves R., Oh S.Y., Liu C., Liu J., Clark A.G., Messer P.W. 2017 Novel CRISPR/Cas9 gene drive constructs reveal insights into mechanisms of resistance allele formation and drive efficiency in genetically diverse populations. PLoS Genet 13(7), e1006796. (doi: 10.1371/journal.pgen.1006796).

11. Marshall J.M., Buchman A., Sanchez C.H., Akbari O.S. 2017 Overcoming evolved resistance to population-suppressing homing-based gene drives. Sci Rep 7(1), 3776. (doi: 10.1038/s41598-017-02744-7).

12. Champer J., Liu J., Oh S.Y., Reeves R., Luthra A., Oakes N., Clark A.G., Messer P.W. 2018 Reducing resistance allele formation in CRISPR gene drive. Proceedings of the National Academy of Sciences 115(21), 5522–5527. (doi: 10.1073/pnas.1720354115).

13. Baker D.A., Nolan T., Fischer B., Pinder A., Crisanti A., Russell S. 2011 A comprehensive gene expression atlas of sex- and tissue-specificity in the malaria vector, Anopheles gambiae. BMC Genomics 12, 296. (doi: 10.1186/1471-2164-12-296).

14. Tazuke S.I., Schulz C., Gilboa L., Fogarty M., Mahowald A.P., Guichet A., Ephrussi A., Wood C.G., Lehmann R., Fuller M.T. 2002 A germline-specific gap junction protein required for survival of differentiating early germ cells. Development 129(10), 2529–2539.

15. Magnusson K., Mendes A.M., Windbichler N., Papathanos P.A., Nolan T., Dottorini T., Rizzi E., Christophides G.K., Crisanti A. 2011 Transcription regulation of sex-biased genes during ontogeny in the malaria vector Anopheles gambiae. PLoS One 6(6), e21572. (doi: 10.1371/journal.pone.0021572).

16. Berleth T., Burri M., Thoma G., Bopp D., Richstein S., Frigerio G., Noll M., Nusslein-Volhard C. 1988 The role of localization of bicoid RNA in organizing the anterior pattern of the Drosophila embryo. EMBO J 7(6), 1749–1756.

17. Schupbach T., Wieschaus E. 1986 Germline autonomy of maternal-effect mutations altering the embryonic body pattern of Drosophila. Dev Biol 113(2), 443–448. (doi: 10.1016/0012-1606(86)90179-x).

18. Wang C., Lehmann R. 1991 Nanos is the localized posterior determinant in Drosophila. Cell 66(4), 637–647. (doi: 10.1016/0092-8674(91)90110-k).

19. Rangan P., DeGennaro M., Jaime-Bustamante K., Coux R.X., Martinho R.G., Lehmann R. 2009 Temporal and spatial control of germ-plasm RNAs. Curr Biol 19(1), 72–77. (doi: 10.1016/j.cub.2008.11.066).

20. Kandul N.P., Liu J., Buchman A., Gantz V.M., Bier E., Akbari O.S. 2020 Assessment of a Split Homing Based Gene Drive for Efficient Knockout of Multiple Genes. G3 (Bethesda) 10(2), 827–837. (doi: 10.1534/g3.119.400985).

21. Akbari O.S., Papathanos P.A., Sandler J.E., Kennedy K., Hay B.A. 2014 Identification of germline transcriptional regulatory elements in Aedes aegypti. Sci Rep 4, 3954. (doi: 10.1038/srep03954).

22. Li M., Yang T., Kandul N.P., Bui M., Gamez S., Raban R., Bennett J., Sanchez C.H., Lanzaro G.C., Schmidt H., et al. 2020 Development of a confinable gene drive system in the human disease vector Aedes aegypti. Elife 9. (doi: 10.7554/eLife.51701).

23. Adelman Z.N., Jasinskiene N., Onal S., Juhn J., Ashikyan A., Salampessy M., MacCauley T., James A.A. 2007 nanos gene control DNA mediates developmentally regulated transposition in the yellow fever mosquito Aedes aegypti. Proc Natl Acad Sci U S A 104(24), 9970–9975. (doi: 10.1073/pnas.0701515104).

24. Macias V.M., Jimenez A.J., Burini-Kojin B., Pledger D., Jasinskiene N., Phong C.H., Chu K., Fazekas A., Martin K., Marinotti O., et al. 2017 nanos-Driven expression of piggyBac transposase induces mobilization of a synthetic autonomous transposon in the malaria vector mosquito, Anopheles stephensi. Insect Biochem Mol Biol 87, 81–89. (doi: 10.1016/j.ibmb.2017.06.014).

25. Meredith J.M., Underhill A., McArthur C.C., Eggleston P. 2013 Next-generation site-directed transgenesis in the malaria vector mosquito Anopheles gambiae: self-docking strains expressing germline-specific phiC31 integrase. PLoS One 8(3), e59264. (doi: 10.1371/journal.pone.0059264).

26. Hong C.C., Hashimoto C. 1995 An unusual mosaic protein with a protease domain, encoded by the nudel gene, is involved in defining embryonic dorsoventral polarity in Drosophila. Cell 82(5), 785–794. (doi: 10.1016/0092-8674(95)90475-1).

27. LeMosy E.K., Hashimoto C. 2000 The nudel protease of Drosophila is required for eggshell biogenesis in addition to embryonic patterning. Dev Biol 217(2), 352–361. (doi: 10.1006/dbio.1999.9562).

28. Galizi R., Hammond A., Kyrou K., Taxiarchi C., Bernardini F., O’Loughlin S.M., Papathanos P.A., Nolan T., Windbichler N., Crisanti A. 2016 A CRISPR-Cas9 sex-ratio distortion system for genetic control. Sci Rep 6, 31139. (doi: 10.1038/srep31139).

29. Champer J., Chung J., Lee Y.L., Liu C., Yang E., Wen Z., Clark A.G., Messer P.W. 2019 Molecular safeguarding of CRISPR gene drive experiments. Elife 8. (doi: 10.7554/eLife.41439).

30. Guichard A., Haque T., Bobik M., Xu X.S., Klanseck C., Kushwah R.B.S., Berni M., Kaduskar B., Gantz V.M., Bier E. 2019 Efficient allelic-drive in Drosophila. Nat Commun 10(1), 1640. (doi: 10.1038/s41467-019-09694-w).

31. Deredec A., Godfray H.C., Burt A. 2011 Requirements for effective malaria control with homing endonuclease genes. Proc Natl Acad Sci U S A 108(43), E874–880. (doi: 10.1073/pnas.1110717108).

32. Legros M., Lloyd A.L., Huang Y., Gould F. 2009 Density-dependent intraspecific competition in the larval stage of Aedes aegypti (Diptera: Culicidae): revisiting the current paradigm. J Med Entomol 46(3), 409–419. (doi: 10.1603/033.046.0301).

33. Beaghton A.K., Hammond A., Nolan T., Crisanti A., Burt A. 2019 Gene drive for population genetic control: non-functional resistance and parental effects. Proc Biol Sci 286(1914), 20191586. (doi: 10.1098/rspb.2019.1586).

34. Deredec A., Burt A., Godfray H.C. 2008 The population genetics of using homing endonuclease genes in vector and pest management. Genetics 179(4), 2013–2026. (doi: 10.1534/genetics.108.089037).

35. Oberhofer G., Ivy T., Hay B.A. 2019 Cleave and Rescue, a novel selfish genetic element and general strategy for gene drive. Proc Natl Acad Sci U S A 116(13), 6250–6259. (doi: 10.1073/pnas.1816928116).

36. Pham T.B., Phong C.H., Bennett J.B., Hwang K., Jasinskiene N., Parker K., Stillinger D., Marshall J.M., Carballar-Lejarazu R., James A.A. 2019 Experimental population modification of the malaria vector mosquito, Anopheles stephensi. PLoS Genet 15(12), e1008440. (doi: 10.1371/journal.pgen.1008440).

37. Giraldo-Calderon G.I., Emrich S.J., MacCallum R.M., Maslen G., Dialynas E., Topalis P., Ho N., Gesing S., VectorBase C., Madey G., et al. 2015 VectorBase: an updated bioinformatics resource for invertebrate vectors and other organisms related with human diseases. Nucleic Acids Res 43(Database issue), D707–713. (doi: 10.1093/nar/gku1117).

38. Fuchs S., Nolan T., Crisanti A. 2013 Mosquito transgenic technologies to reduce Plasmodium transmission. Methods Mol Biol 923, 601–622. (doi: 10.1007/978-1-62703-026-7_41).

39. Volohonsky G., Terenzi O., Soichot J., Naujoks D.A., Nolan T., Windbichler N., Kapps D., Smidler A.L., Vittu A., Costa G., et al. 2015 Tools for Anopheles gambiae Transgenesis. G3 (Bethesda) 5(6), 1151–1163. (doi: 10.1534/g3.115.016808).

40. Pinello L., Canver M.C., Hoban M.D., Orkin S.H., Kohn D.B., Bauer D.E., Yuan G.C. 2016 Analyzing CRISPR genome-editing experiments with CRISPResso. Nat Biotechnol 34(7), 695–697. (doi: 10.1038/nbt.3583).

## Supplementary References

[1] Papathanos, P. A., Windbichler, N., Menichelli, M., Burt, A. and Crisanti, A. The vasa regulatory region mediates germline expression and maternal transmission of proteins in the malaria mosquito Anopheles gambiae: a versatile tool for genetic control strategies. BMC Mol Biol 10, 65, (2009).

[2] Hammond, A.M. et al. The creation and selection of mutations resistant to a gene drive over multiple generations in the malaria mosquito. PLoS Genet 13, e1007039 (2017).

[3] Kyrou K., Hammond A.M., Galizi R., Kranjc N., Burt A., Beaghton A.K., Nolan T., Crisanti A. 2018 A CRISPR-Cas9 gene drive targeting doublesex causes complete population suppression in caged Anopheles gambiae mosquitoes. Nat Biotechnol 36(11), 1062–1066. (doi:10.1038/nbt.4245)

[4] Beaghton A.K., Hammond A., Nolan T., Crisanti A., Burt A. 2019 Gene drive for population genetic control: non-functional resistance and parental effects. Proc Biol Sci 286 (1914), 20191586. (doi:10.1098/rspb.2019.1586)

[5] Wolfram Research, Inc., 2017 Mathematica 11.2, Champaign, IL.

